# Physiologically Based Multiphysics Pharmacokinetic Model for Determining the Temporal Biodistribution of Targeted Nanoparticles

**DOI:** 10.1101/2022.03.07.483218

**Authors:** Emma Glass, Sahil Kulkarni, Christina Eng, Shurui Feng, Avishi Malavia, Ravi Radhakrishnan

## Abstract

Nanoparticles (**NP**) are being increasingly explored as vehicles for targeted drug delivery because they can overcome free therapeutic limitations by drug encapsulation, thereby increasing solubility and transport across cell membranes. However, a translational gap exists from animal to human studies resulting in only several NP having FDA approval. Because of this, researchers have begun to turn toward physiologically based pharmacokinetic (**PBPK**) models to guide in vivo NP experimentation. However, typical PBPK models use an empirically derived framework that cannot be universally applied to varying NP constructs and experimental settings. The purpose of this study was to develop a physics-based multiscale PBPK compartmental model for determining continuous NP biodistribution. We successfully developed two versions of a physics-based compartmental model, models A and B, and validated the models with experimental data. The more physiologically relevant model (model B) had an output that more closely resembled experimental data as determined by normalized root mean squared deviation (NRMSD) analysis. A branched model was developed to enable the model to account for varying NP sizes. With the help of the branched model, we were able to show that branching in vasculature causes enhanced uptake of NP in the organ tissue. The models were solved using two of the most popular computational platforms, MATLAB and Julia. Our experimentation with the two suggests the highly optimized ODE solver package DifferentialEquations.jl in Julia outperforms MATLAB when solving a stiff system of ordinary differential equations (**ODEs**). We experimented with solving our PBPK model with a neural network using Julia’s Flux.jl package. We were able to demonstrate that a neural network can learn to solve a system of ODEs when the system can be made non-stiff via quasi-steady-state approximation (**QSSA**). In the future, this model will incorporate modules that account for varying NP surface chemistries, multiscale vascular hydrodynamic effects, and effects of the immune system to create a more comprehensive and modular model for predicting NP biodistribution in a variety of NP constructs.

**Author summary:** Nanoparticles (NP) have been used in various drug delivery contexts because they can target specific locations in the body. However, there is a translational gap between animals and humans, so researchers have begun toward computational models to guide in vivo NP experimentation. Here, we present several versions of physics-based multiscale physiologically based pharmacokinetic models (PBPK) for determining NP biodistribution. We successfully developed two versions of ODE-based compartmental models (models A and B) and an ODE-based branched vascular model implemented in MATLAB and Julia and validated models with experimental data. Additionally, we demonstrated using a neural network to solve our ODE system. In the future, this model can integrate different NP surface chemistries, immune system effects, multiscale vascular hydrodynamic effects, which will enhance the ability of this model to guide a variety of in vivo experiments.

## Introduction

Clinical medicine has entered an era where nanotechnology is becoming more and more prevalent, with the use of drug-carrying nanoparticles (**NP**) increasing in recent years. NP can overcome free therapeutic limitations by drug encapsulation, thereby enhancing solubility by promoting transport across cellular membranes [1]. Because of this, NPs are increasingly being explored as vehicles for targeted drug delivery to healthy and cancerous tissues, for diagnostic imaging purposes, and to enhance T-cell-based immunotherapies ([2], [3]). While extensive studies are being performed to determine the efficacy of certain NP based therapeutics in *in vitro* and *in vivo* animal models, there exists a translational gap from animal to human studies resulting in only a select few NP being approved for FDA use [4].

One major underlying cause of this translational gap is the difference in the physiology of animal models compared to humans, which have the potential to affect NP behavior and functionality [5]. However, the cause of the translational gap that is the motivation for this study is the heterogeneity of NP constructs and experimental models. There are nearly endless NP constructs (e.g., rigid, flexible, spherical, non-spherical, polymeric, DNA-based, etc.), sizes (a few nm to a few microns), and experimental models for translational studies, making NP a complex mix of biology and engineering. The wide array of applications, targets, and physical characteristics of NP significantly impedes the ability of NP to be researched effectively as possible bench-to-bedside therapeutics.

To this end, researchers have begun to turn toward physiologically based pharmacokinetic (**PBPK**) models to guide in vivo experimentation and better understand NP targeting behavior and performance in the human body, resulting in more effective and efficient usage for a variety of previously described applications ([6], [8], [9], [10]). Traditional pharmacokinetic (PK) models describe the concentration of drugs in the blood plasma over time ([11], [12]), while PBPK models consist of compartments that represent actual tissues and organ spaces. Existing PBPK models for small molecules or biologics ([13], [14], [15]) use an empirically derived framework for parameterization, resulting in a model that cannot be universally applied with varying NP constructs and experimental settings ([16], [17]). Additionally, multiphysics aspects, including physiological and hydrodynamic factors governing NP biodistribution and tissue targeting, involve mechanisms operating at multiple lengths and timescales ([18]). Therefore, a multiscale computationally driven model with physiologically relevant inputs can be utilized to understand the organ-specific biodistribution of NP. The multiscale model must incorporate NP hydrodynamic properties in the vasculature, NP-Endothelial Cell (**EC**) adhesion properties, and NP subcellular interactions that govern targeted uptake to fully describe the movement and accumulation of NP within the body [6]. In order to create a comprehensive multiscale model, NP behavior must be understood at the system, hydrodynamic, and cell adhesion scales.

A previously published multiscale PBPK model has determined binding constants of intracellular adhesion molecule 1 (**ICAM1**) coated NPs to endothelial cell (EC) surface receptors in mice and humans by utilizing the biophysical properties of the antibody to receptor interactions, and the cell surface [16]. Additionally, this model determines the percent injected dose (**%IDG**) that distributes to the tissue in given organs at steady-state; non-specific uptake is not accounted for in this model (NP uptake via passive diffusion in the intercellular cleft). The binding constants determined through the previous model [16] model will help to drive the binding characteristics of NP in the multiscale model described in this study. Additionally, the rate at which NP bound to the EC layer are endocytosed into a specific organ tissue must be considered. The concentration of NP retained within the tissue or biodistribution is ultimately most important since this allows for understanding how effectively NP can target tissue.

The purpose of this study is to 1) advance existing steady-state multiscale PBPK models [16] to incorporate NP uptake via nonspecific transport, 2) develop novel multiscale PBPK compartmental models to predict temporal effects, and 3) introduce a compartmental branched vascular model that can predict the effect of hydrodynamic interactions that depend on NP size, flow, and vasculature network properties, 4) perform validation with experimental murine biodistribution data, 5) propose efficient solvers for coupled stiff systems that embody the above properties, as well as make the solvers compatible with contemporary machine-learning-based modules (such as neural networks) which can capture and incorporate multiphysics models in the PBPK workflow.

## Materials and methods

### Overview

Many Monte Carlo-based models have been proposed in previous works and have been integral in understanding multivalent receptor-NP interactions at a molecular level. Agrawal and Radhakrishnan [19] quantitatively characterized NP-endothelial cell interactions by determining the multivalence of NP binding as well as antigen clustering, ultimately providing future models with details about the energetics of the NP binding process. Ramakrishnan [16] used these binding properties to understand the role that expression levels of NP receptors play when targeting live cells, validating with experimental data. While these adhesion-centric models are necessary for understanding NP characteristics at the molecular level, they do not always include the interaction of NP with the vascular network and translate into the pharmacodynamic scales. By considering hydrodynamic parameters of the vascular system, and cell-binding/uptake parameters into a variety of organ tissues (as determined in previous studies such as [16]), a model that is governed by multiple time scales can be created, providing a more physiologically relevant determination of NP biodistribution with temporal resolution. This paper describes three variations of a multiscale pharmacokinetic model: 1) a steady-state model that facilitates the translation of the multivalent free energy of adhesion into a biodistribution; 2) two different compartmental models (models A and B) that are physiologically based and can predict temporal biodistribution; and 3) a compartmental model that incorporates the vascular branching network and can include multiphysics effects such as hydrodynamic interactions, NP size-dependent effects of flow and adhesion, with temporal resolution [6, 7]. We describe the steady-state or continuous temporal biodistribution of ICAM-1 targeted NP for a murine model in each case. The multiscale nature of the model described here can allow for customization with various NP sizes, shapes, and uptake parameters, resulting in a customizable predictive platform for different NP chemistries, organisms, and pathophysiologies.

### Steady-State Model

A steady-state, multiscale, PBPK model was developed by Ramakrishnan [16] to determine the percent injected dose of NP per gram of tissue (%idg) in five organ compartments: lung, heart, kidney, liver, and spleen. However, this model only considers NP uptake via antibody-receptor mediated multivalent interactions, failing to account for receptor-mediated internalization and non-specific transport through pores between the intercellular cleft of adjacent endothelial cells, especially in clearance organs. Here, the existing steady-state PBPK model was modified to incorporate this non-specific uptake, to have the modified model better predict in vivo biodistribution data at short times (*<* 30 min, when NP does not internalize substantially) than the original model as determined by the R (square root of the coefficient of determination R^2^) values. Several versions of the model (incorporating different multiphysics) are considered based on specific adhesion of NP to epithelial cell membrane surfaces, including flat membrane and membrane mimicking live cells. In addition, the presence of resident macrophages and activated macrophages are considered.

The original model [16] is described using Eq (1),

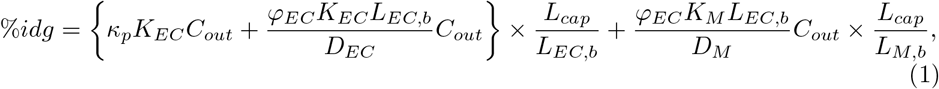

where *κ*_*p*_ is the non-specific binding of NP, *K*_*EC*_ is the association constant for binding of NP to endothelial cell surface receptors, *K*_*M*_ is the association constant for NP binding to a macrophage cell, *D*_*EC*_ and *D*_*M*_ represent the diameter of endothelial cells and macrophage cells respectively, *φ*_*M*_ and *φ*_*EC*_ represent the concentration of endothelial cells and macrophage cells in the target tissue respectively, *L*_*EC,b*_ and *L*_*M,b*_ represents the distance from an EC or macrophage surface receptor that the NP can successfully bind respectively, *C*_*out*_ represents the injected concentration of NP, and *L*_*cap*_ represents the size of the cell-free layer in the capillary in which the NP is perfused. Incorporating the non-specific transport of NP into the tissue, the modified model equation can be described using Eq (2):

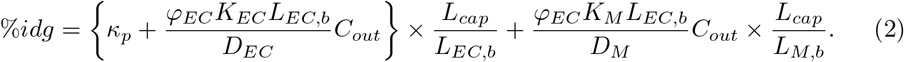

The model above is expected to approximate the biodistribution at short timescales, defined as the regime in which the observation time is less than the timescale for internalization. The model is easy to compute as it does not involve solving dynamics equations owing to its steady-state nature. Later, we show that the temporal model predicts the same behavior as the steady-state model at these short timescales, thereby validating the steady-state approximation utilized here for short time scales.

### Temporal Model

#### Compartmental Model Development

The goal of the compartmental model is to develop a basic framework for determining targeted NP biodistribution in a murine model that can act as a predictive model when provided with experimentally and empirically derived parameters. This model consists of five to seven organ compartments (lung, heart, kidney, liver, spleen and in model A; lung, heart, kidney, liver, spleen, gut, and ‘other in model B) each of which is interconnected via the arterial and venous compartments. The ‘other’ compartment consists of all organs that are not explicitly included in the model. We have developed two ways in which the organ compartments can be connected by the arteries and veins, shown in Figure 1. Figure 1a, is the model A configuration (an oversimplified version of the circulatory system with 5 organ compartments), while Figure 1b shows the model B configuration model, which is more physiologically relevant (lung circulation has been separated out to keep track of oxygenation and oxygen distribution, and gut and spleen compartments are coupled to the liver compartment, and gut/other compartments were incorporated as well).

**Fig 1.**
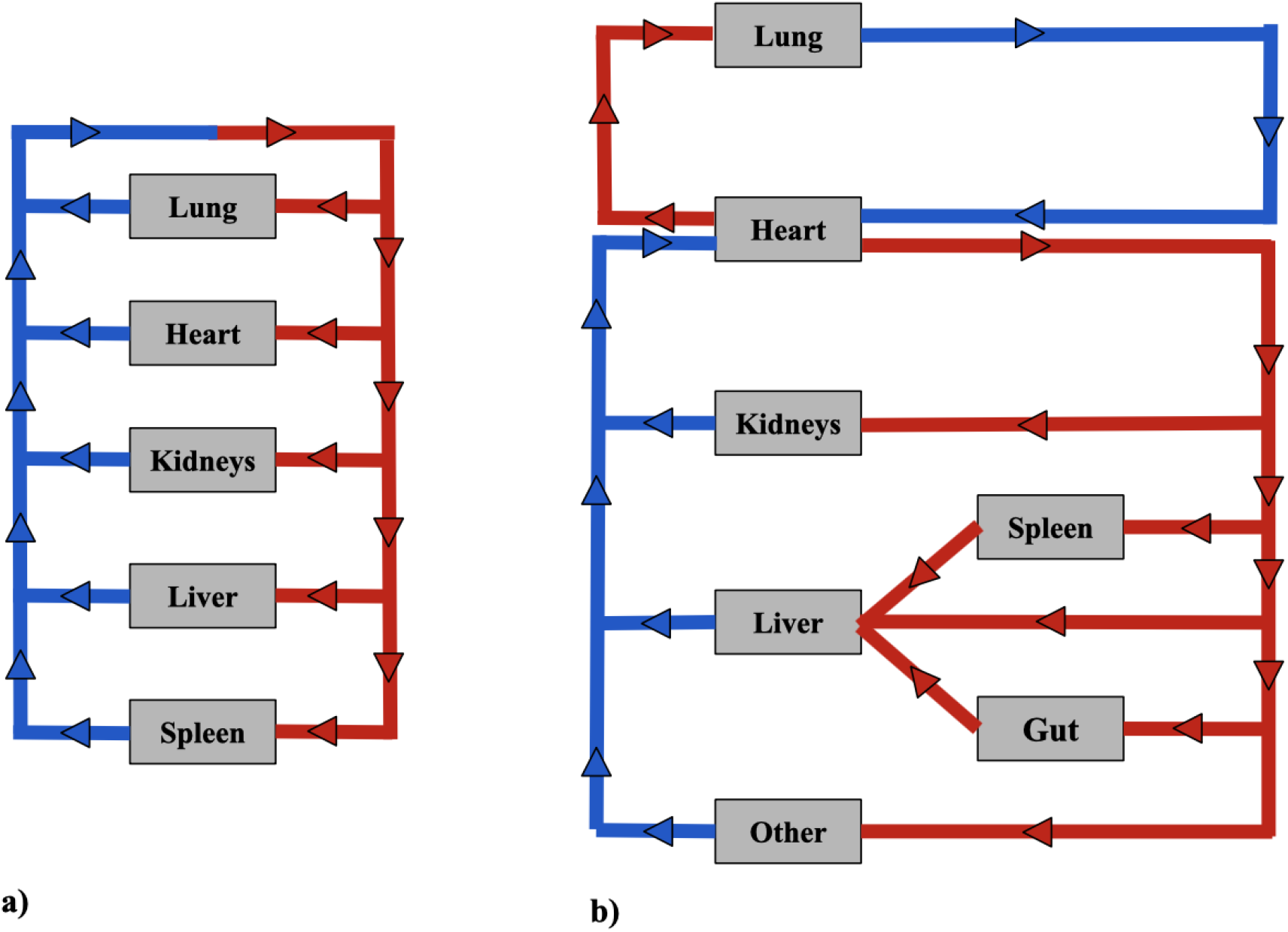
Compartmental model configurations. a) model A configuration. b) model B configuration.

Each organ compartment in either model configuration consists of three additional compartments: vascular, endothelial cell, and tissue compartments shown in Figure 2. As NPs enter an organ compartment through the vascular compartment, they can either bind to the ICAM-1 endothelial cell surface receptors (*K*_*on*_), enter the organ tissue compartment via non-specific uptake (*K*_*NS*_), or be degraded (*K*_*deg*_). Then, once bound to the endothelial cell layer, the NP can either unbind from the EC compartment returning to the vascular compartment (*K*_*off*_), be taken into the organ tissue via transcytosis (*K*_*up*_), or degraded (*K*_*deg*_). Once NPs are in the tissue compartment, they can be degraded (*K*_*deg*_). It is also important to note that NP can be degraded (*K*_*deg*_) within the arterial and venous compartments as well. The difference in time scales represented in the arteries/veins and the cellular scale compartments contribute to the temporal multiscale nature of the model.

**Fig 2.**
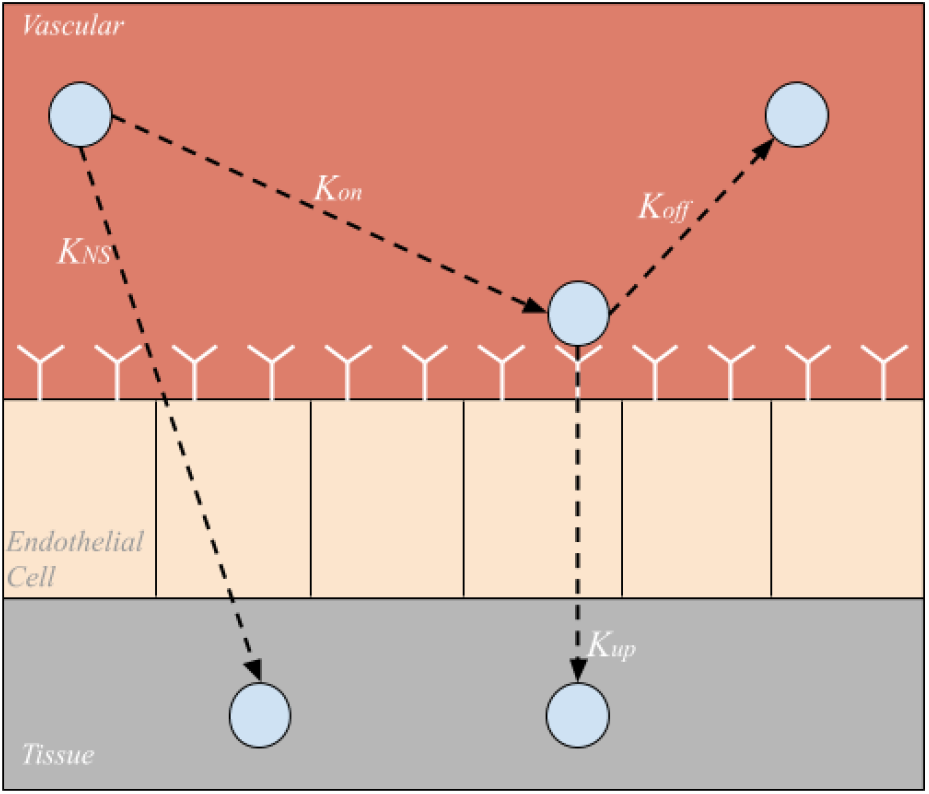
Sub-compartments within each organ compartment. Nanoparticles can be taken directly into the tissue via nonspecific uptake (*K*_*NS*_), bound to endothelial cell surface receptors (*K*_*on*_), unbound from endothelial cell surface receptors (*K*_*off*_), or taken into the tissue via endo or transcytosis (*K*_*up*_).

#### Model A Equations

The model A configuration can be described using a system of 17 linear ordinary differential equations (ODEs). Model A is described using equations 3-7, where *i* = lung, heart, kidneys, liver, spleen. The venous compartment is described by:

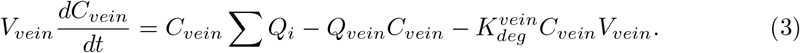

Eq (3) describes the change in concentration of NP in the venous compartment over time, where *Q*_*i*_ and *Q*_*vein*_ represent the flow of blood through organ compartments and the veins, respectively, *V*_*vein*_ denotes the volume of blood in the vein, *C*_*vein*_ is the concentration of NP in the veins, and 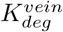 is the degradation rate of NP in the veins. The arterial compartment is described by:

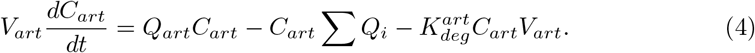

Eq (4) describes the change in concentration of NP in the arterial compartment over time, where *Q*_*art*_ represents the flow of blood through the arteries, *V*_*art*_ denotes the volume of blood in the arteries, *C*_*art*_ is the concentration of NP in the arteries, and 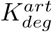 is the degradation rate of NP in the arteries. For each tissue type *i*, the vascular compartment is described by:

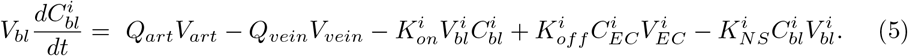

Eq (5) describes the change in concentration of NP in the vascular compartment within each organ compartment of the model over time, where 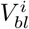 and 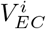 represent the volume of blood in the vascular and the endothelial cell compartments (which is defined by the product of the length of the endothelial cell receptors and the surface area of the vascular compartment) of the organ respectively, 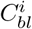 and 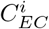 denotes the concentration of NP in the vascular and endothelial cell compartments of the organ respectively, 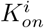 denotes the rate of binding of NP to the endothelial cell surface receptors, 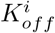 denotes the rate of NP unbinding from the endothelial cell surface receptors, and 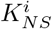 denotes the rate of nonspecific NP uptake into the organ tissue. The endothelial compartment in each tissue is described by:

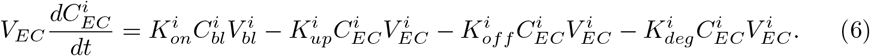

Eq (6) describes the change in concentration of NP bound to the endothelial cell surface receptors over time, where 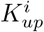 denotes the rate of uptake of NP into the organ tissue via transcytosis. Each tissue compartment is described by:

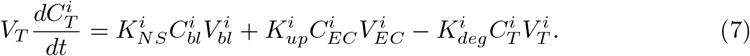

Eq (7) describes the change in concentration of NP in the organ tissue over time where 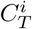 and 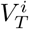 denote the concentration of NP in the tissue compartment of the organ and the volume of the tissue compartment respectively.

#### Model B Equations

The model B configuration (Figure 1b) can be described using a system of 23 linear ODEs, however, due to the addition of two organs and the alternative configuration of the model, several equations differ, while the overall structure is the same. Equations 8-13 Describe the modified model configuration:

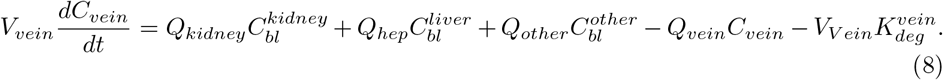

Eq (8) describes the change in concentration of NP in the veins over time, where *Q*_*kidney*_ and *Q*_*other*_ denote the flow rate of blood through the kidneys and ‘other’ compartment respectively, *Q*_*hep*_ denotes the combined flow rate of blood through the liver, spleen, and gut compartments (*Q*_*hep*_ = *Q*_*liver*_ + *Q*_*spleen*_ + *Q*_*gut*_). *C*_*bl*_ denotes the concentration of NP in the vascular compartment of each respective organ (lung, heart, kidneys, liver, spleen, gut, other)

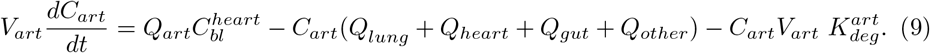

Eq (9) describes the change in concentration of NP in the arteries over time where *Q*_*lung*_ and *Q*_*heart*_ denote the flow rate of blood through the lung and heart compartments, and 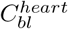 is the concentration of NP in the vascular compartment of the heart.

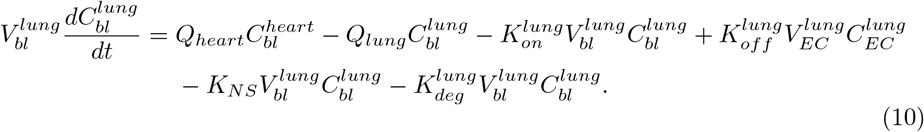

Eq (10) describes the change in concentration of NP in the vascular compartment of the lung over time.

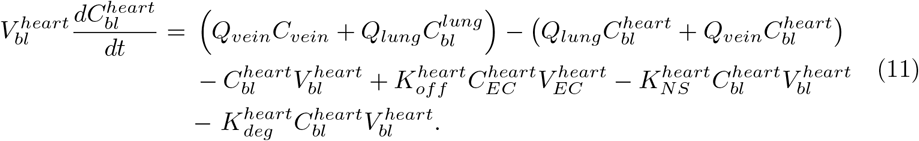

Eq (11) describes the change in concentration of NP in the vascular compartment of the heart over time.

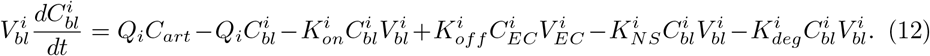

Eq (12) describes the change in concentration of the NP in the vascular compartment of the kidneys, spleen, gut, and ‘other’ compartment over time (where i = kidney, spleen, gut, other).

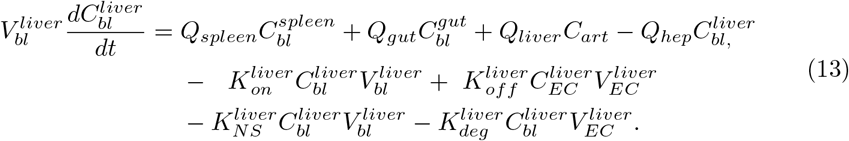

Eq (13) describes the change in concentration of the NP in the vascular compartment of the liver over time. Eq (6) and Eq (7) were used in the original model to describe the change in concentration of NP in the endothelial cell compartments and tissue compartments respectively, can also be used in the modified model configuration.

Additionally, Eq (14) was used to express the total NP degraded in the system at a given time *t*, which will be useful to determine mass conservation of the system.

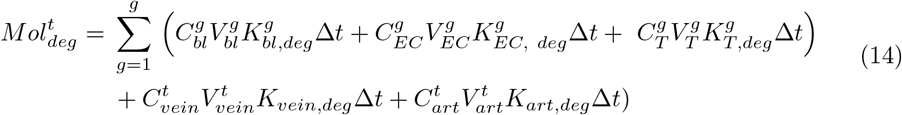

where *t* is the total time the model is run, g is the total number of organs in the model, and Δ*t* is the *t* step of the model.

Eq (15) was used to determine the mass conservation of the system. If the system is closed, the line formed by 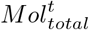 over every time point *t* of the simulation should have a slope of 0.

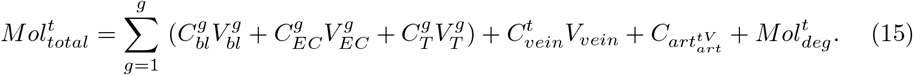

Eq (16) describes the mass conservation in molar basis, this equation was used to determine the mass conservation when the system of ODE was set up in terms of molar profiles.

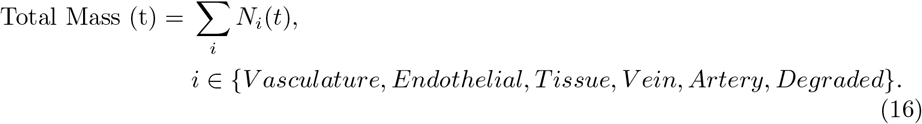

### Vasculature Branching

#### Branched Model Development

The purpose of developing a branched model was to create a more detailed and physiologically relevant version of the basic compartmental model. Additionally, utilizing this branched model will ultimately increase the specificity of uptake rate constants for NP of various sizes. Furthermore, the branched model will enable the inclusion of key margination and hydrodynamic interactions whose effects are determined by the flow rate and blood hematocrit concentration. The branched network consists of a branched vascular tree that begins at the main arteries and veins and bifurcates into the capillary beds, connecting the arterial and venous branching networks (Figure 3). While asymmetric branching patterns characterize typical vasculature networks, for simplicity, the branching model described here will consist of identical daughter vessel segments at each generation of bifurcation. Daughter vessels will continue to bifurcate, their diameters following the power-law relationship until the diameter of the vessel approaches the size of a red blood cell (the smallest vessel in our network). Below the development of the branched network is discussed in greater detail.

**Fig 3.**
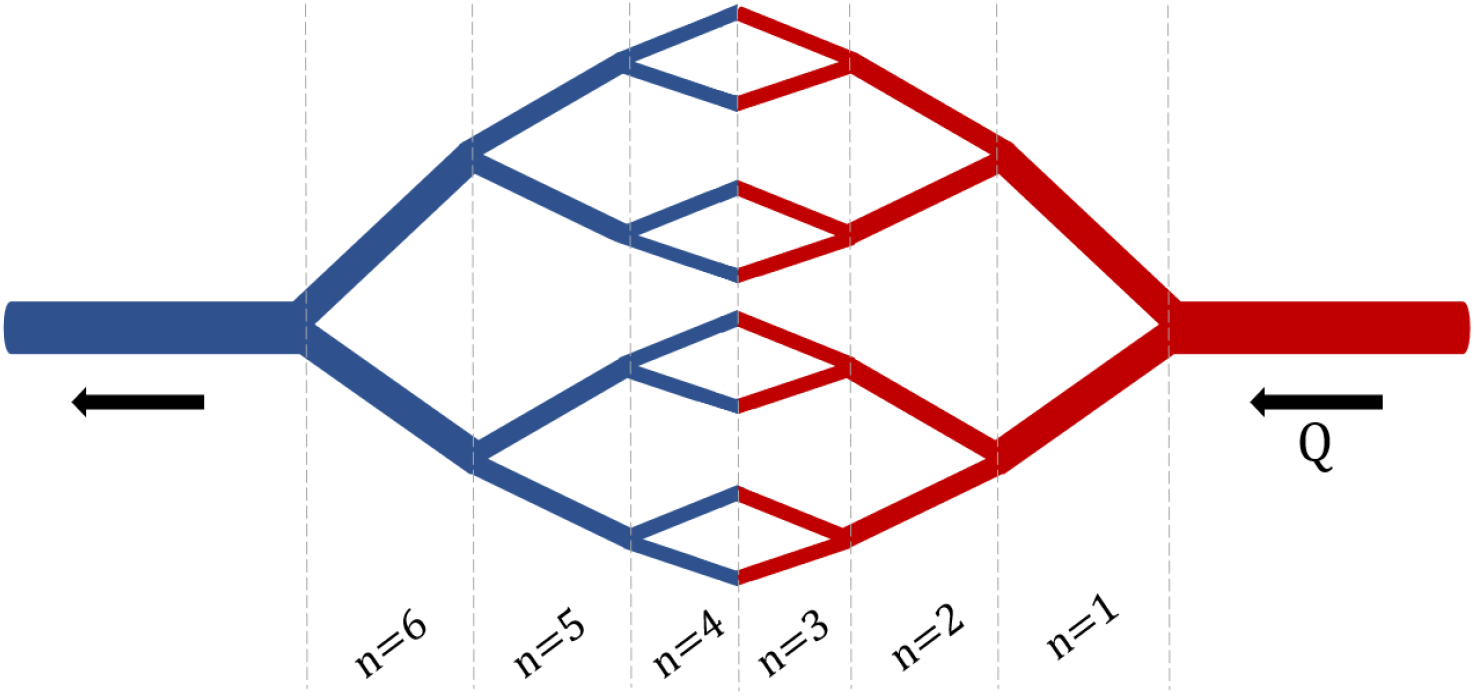
Branched Network. An example branching network with *i* = 3 bifurcations and *n* = 6 generations.

The development of this branched network was modeled after the branching network described in [20], but is a simplified version of their three-dimensional vascular branching network model. The diameter of the daughter vessels is governed by the power law relationship, where the parent vessel is *d*_0_ and daughter vessels are *d*_1_ and *d*_2_, and *k*=3, which is typically assumed based on a minimum dissipation principle representing a stable flow condition:

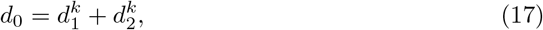

where *d*_0_ *> d*_1_ = *d*_2_

Due to the branching nature of the model, we can determine the number of segments (*N*) at any generation of branching (*n*) using the relationship *N* = 2^*n*^. While the number of segments increases as the number of generations increase, the diameter of the branches decreases. Suppose that the diameter of the parent vessel at a given bifurcation (*i*) is *d*_*i*,0_, and the diameter of the daughter vessels are *d*_*i*,1_ and *d*_*i*,2_. Given a symmetric bifurcation (*d*_*i*,1_ = *d*_*i*,2_), the following relationship can be used to relate the diameter of the parent vessels to that of the daughter vessels.

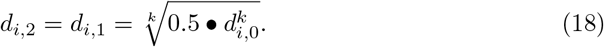

Then, the length of each vessel can be determined using the know length to diameter ratio *β* = 3, and the diameter of the vessel, *d*.

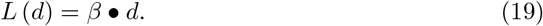

This network will begin at the diameter of the main artery *d*_0_ = 600 *μ* m (main artery diameter in mouse), continue to bifurcate until the diameter of the daughter vessels reaches the diameter of a red blood cell (*d* = 15 *μ* m). This lower-bound in vessel diameter results in a network of 16 bifurcations and 32 generations of branching for each branching element. A schematic of this branching network can be seen in Figure 3.

After constructing this branched network, it is important to determine the surface area and volume of the vascular network element as a whole. Supposing *d*_*i*_, *l*_*i*_, and *N*_*i*_ are the diameter, length, and the number of daughter vessels in a given generation *i* respectively, Equation (20) can be used to describe the total vessel surface area in a branching element.

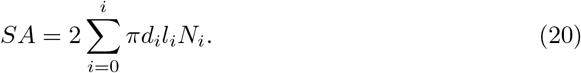

Equation (21) can be used to determine the total vascular volume of a branching element,

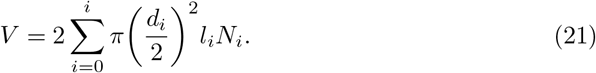

It is important to note that the number of generations, volume, and surface area of one branching element is fixed. I.e., the volume of a single branching element does not differ across organs. Each organ will have a different number of branching elements dependent on *ϕ*_*i*_, which is the ratio of total organ vascular volume to that of one branching element.

#### 0.0.1 Branched Model Equations

The system of ODEs used to describe the branched compartmental model consists of 457 equations. The organs are organized in the model B configuration, and therefore the equations in the branched model take on a similar form when describing the transport between organs. Therefore, the equations describing the concentration of NP in the arteries and veins over time are the same as in the modified compartmental model configuration, Equations (8) and (9), for veins and arteries, respectively.

Eqs 21-32 describe the rest of the branched vascular compartmental model that bifurcates from the diameter of the main vein to the size of a red blood cell and branches back out to the diameter of the main artery. Each organ compartment contains four types of equations to describe the concentration of NP in the vasculature: 1) one equation describing NP concentration in the first generation (*n* = 1), 2) series of equations describing NP concentration in the branching network until the diameter of the branch is that of a red blood cell (*n* = 2 to *n* = 16), 3) series of equations describing NP concentration in branching network until diameter of branch branches back out to the generation before the diameter of the vasculature is that of the main artery (*n* = 17 to *n* =31), 4) one equation describing the NP concentration in the last generation (*n* = 32).

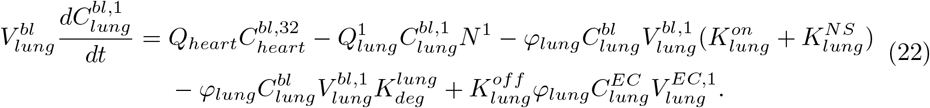

Eq (22) describes the NP concentration in generation *n* = 1 in the vascular branching network of the lung. 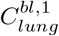 denotes the NP concentration of the lung vasculature in generation *n* = 1 of the branching network, 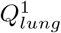 represents the flow rate through generation *n* = 1 of the lung vascular network, 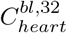 denotes the NP concentration of the heart vasculature in generation *n* = 32 (the concentration of NP exiting the heart compartment), *N* ^1^ represents the number of branches in generation *n* = 1, *φ*_*lung*_ represents the total number of branching elements in the given organ compartment, 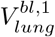 denotes the vascular volume of a single branching element in generation *n* = 1, and 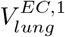 denotes the volume of the lung endothelial cells of a single branching element in generation *n* = 1.

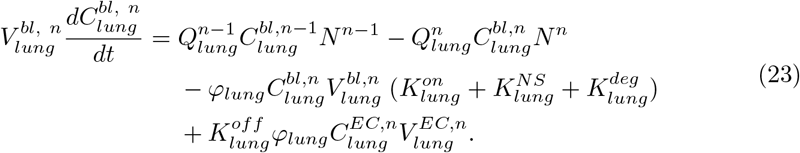

Eq (23) describes the NP concentration in the lung vasculature in generation *n* = 2 to *n* = 16 of the branching network. 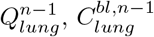, and *N* ^*n*−1^ denote the flow rate, NP concentration, and number of branches in the lung vasculature of the previous generation of the branching network.

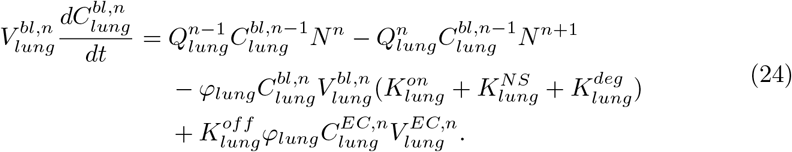

Equation (24) describes the NP concentration in the lung vasculature in generation *n* = 17 to *n* = 31 of the branching network.

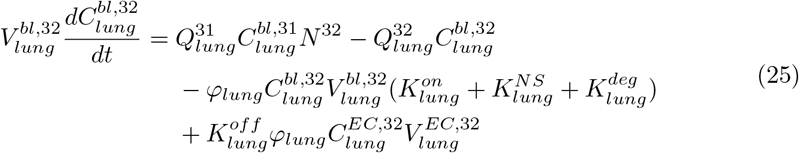

Equation (25) describes the NP concentration in the lung vasculature in generation *n* = 32 of the branching network. The heart compartment vascular branching network can be described similarly to the lung compartment, with a series of four equations.

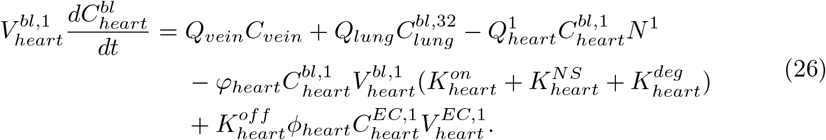

Equation (26) describes the NP concentration in the heart vasculature in generation *n* = 1 of the branching network. The equations describing the NP concentration in the heart vasculature in generations *n* = 2 to *n* = 16 and *n* = 17 to *n* = 31 are the same as equations (23) and (24) respectively, but *i* = lung should be replaced with *i =* heart.

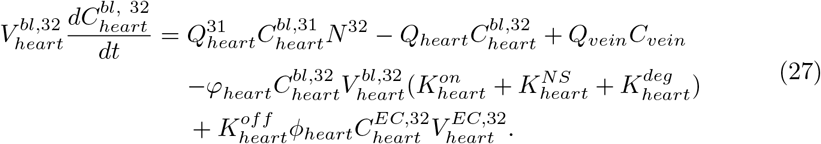

Equation (27) describes the NP concentration in the heart vasculature in generation *n* = 32 of the branching network. The equations describing the concentration of NP in the liver compartment are again constructed similarly to the branching equations in the lung and heart compartments and again consist of a series of four equations.

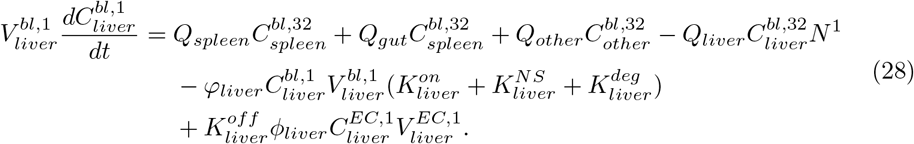

Equation (28) describes the NP concentration in the liver vasculature in generation *n* = 1 of the branching network. The equations describing the NP concentration in the liver vasculature in generations *n* = 2 to *n* = 16 and *n* = 17 to *n* = 31 are the same as equations (23) and (24) respectively, but *i* = lung should be replaced with *i =* liver.

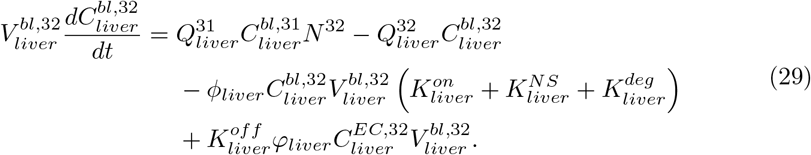

Equation (29) describes the NP concentration in the liver vasculature in generation *n* = 32 of the branching network. The equation describing the NP concentration in the kidneys, spleen, gut, and other compartments are again constructed similarly to the branching equations in the lung and heart compartments, and again consist of a series of four equations.

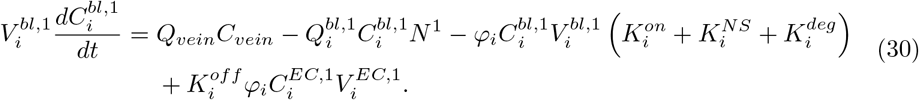

Equation (30) describes the NP concentration in the *i =* kidneys, spleen, gut, and other compartment’s vasculature in generation *n* = 1 of the branching network. The equations describing the NP concentration in the liver vasculature in generations *n* = 2 to *n* = 16 and *n* = 17 to *n* = 31 are the same as equations (23) and (24) respectively, but *i* = lung should be replaced with *i =* kidneys, spleen, gut, other.

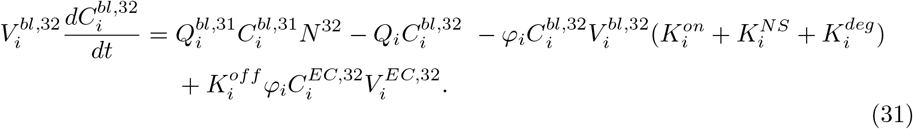

Equation (31) describes the NP concentration in the liver vasculature in generation *n* = 32 of the branching network. Next, we can describe the NP concentration that is bound to the endothelial cell layer.

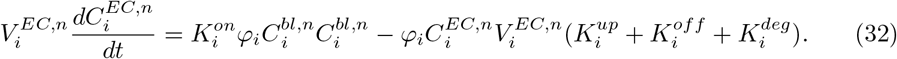

Equation (32) describes the NP concentration bound to the endothelial cells of *i* = lung, heart, kidney, liver, spleen, gut and other in generation *n =*1 to *n* = 32. Finally we can describe the concentration of NP that is distributed to the organ tissue.

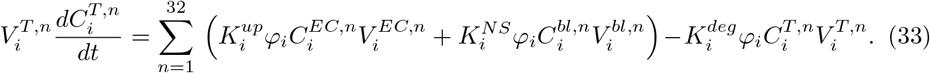

Equation (33) describes the NP concentration in the tissue of *i* = lung, heart, kidneys, liver, spleen, gut, and other in generation *n =* 1 to *n* = 32, where the sum of the antibody-receptor mediated endocytosis 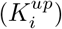 and non-specific uptake 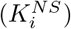 is summed over all generations of branching.

### Parameterization

#### Parameters for Compartmental Model

The compartmental model is parameterized with a variety of physiological inputs such as blood, tissue, endothelial cell volumes, and blood flow rates collected from previously published sources ([21], [22]). Other parameters such as *K*_*on*_ and *K*_*off*_ for ICAM antibody-coated NP were computationally determined in a previous study [16], while yet other rate constants were determined via local sensitivity analysis through comparison to existing translational studies [23].

The flow rates through each individual organ, *Q*_*i*_, were determined using Table 3 (in supporting information) from [21], which gave blood flow rates through mouse organ vasculature in units of L/g/min, which were ultimately converted into units of L/min for use in the compartmental models by using mouse organ weight data from [22] and the female and male masses were averaged. Additionally, to calculate the total flow, *Q*, the sum of flow rates across all organs was taken.

Table 1 (in supporting information) in [21] listed values for vascular volume in various mouse tissues. The volumes were listed in units of L/g. The mouse organ weight data from [22] was used to determine the tissue volume for the entire organ, 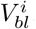.

The volume of blood in the mouse vein was computed using the following equation, 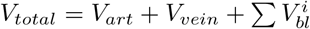, where *V*_*total*_ = 2146 *μ*L. The volume of blood in the veins (*V*_*vein*_) and the arteries (*V*_*art*_) is considered to be the same. Then the volume of blood in the mouse venous system was computed.

Values for interstitial tissue volume and extracellular tissue volume were obtained using Table 2 (in supporting information) from [21]. Values for interstitial tissue volume (given in L/g) were used to determine the volume of each organ in its entirety, 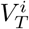. By using the organ weights from [21], as was done to determine flow rates (*Q*_*i*_), the tissue volume for each organ in the mouse was computed with units of L.

The parameter *l*_*NC,b*_ denotes the height of the endothelial cell layer, is constant between organs and species and has already been defined by a previous model [16]. *A*_*i*_ can be calculated by using known relationships from [16], eg, It is known that 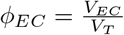 and 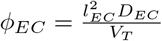. Given *ϕ*= 0.3, *D*_*EC*_ = 5 × 10^−6^, and the *V*_*T*_ for each mouse organ previously described, 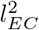 can be calculated. When *A*_*i*_ and *l*_*NC,b*_ are multiplied together, the resulting value will be representative of the volume of the endothelial cell layer; 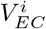.

The binding rate of NP to the endothelial cell surface 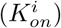 was determined using the relationship 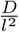 given in [16]. The unbinding rate of NP from the endothelial cell surface 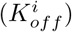 can be determined by using 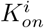 and *K*_*EC*_ from [16] for each organ). However, the log of the *K*_*EC*_ had to be taken to accommodate for crowding effects [24]. So, 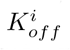 is calculated as a ratio of 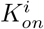 to 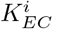; i.e., we assume a diffusion limited on-rate, where the diffusion occurs through the glycocalyx. The off-rate is computed based on the on-rate and the equilibrium constant.

#### Parameterization of NP-Size Dependent Branched Model

The relation between *K*_*EC*_ and particle diameter was obtained from Figure 9 of [24]. A linear relationship between log(*K*_*eq*_)) and particle diameter (a) was established using the slope-point form for every organ, the point on the line being the (a, log(*K*_*EC*_)), for a = 800nm, which was obtained using Monte Carlo simulation in [16]. The binding rate is computed as; 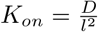 where *l* = thickness of the glycocalyx layer and *D* is given by the Stokes–Einstein equation; 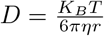, then 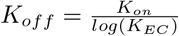. The values of *K*_*on*_, *K*_*off*_, and *log*(*K*_*EC*_), are reported in Table 1, 2, and 3, in supporting information.

#### Sensitivity Analysis

Parameters that do not have an explicit value stated in previously published literature were subject to local sensitivity analysis. These parameters included 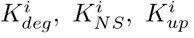; where *i =* lung, heart, kidneys, liver, spleen, gut, and other. To perform the local sensitivity analysis the values of 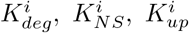 were changed incrementally so the output curve closely resembled that of an experimental data set (discussed in section 2.5.1). Larger 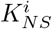 and 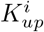 values resulted in a steeper initial biodistribution curve. Generally, 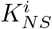 and 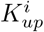 had a similar magnitude result in the biodistribution curve. Larger 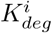 resulted in a smaller maximum biodistribution and a quicker decay in the biodistribution curve. The results of an example local sensitivity analysis for the spleen is shown in Figure 4. The parameters determined via the local sensitivity analysis that were used in the compartmental models can be found at the SI GitHub link. While a local sensitivity analysis was performed on certain parameters, it is important to note that no extensive linear optimization or global sensitivity analysis was done on these parameters; this will be done in future iterations of this work.

**Fig 4.**
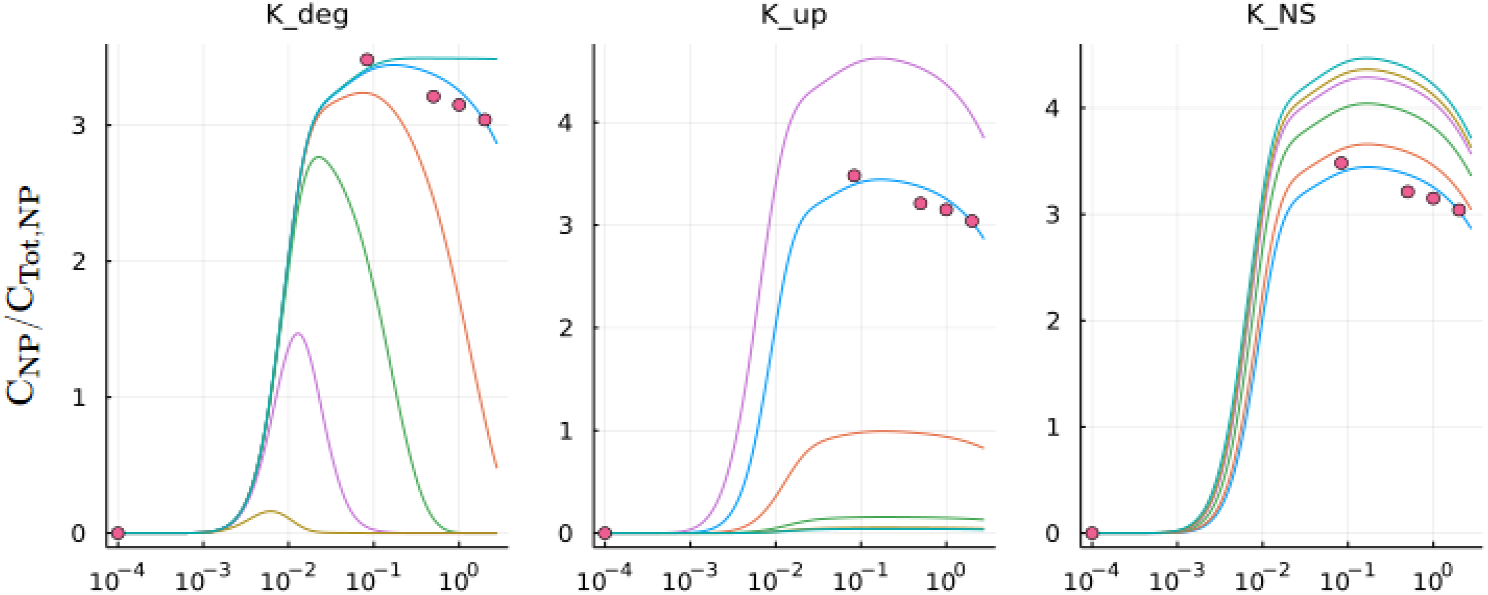
Local sensitivity analysis: Plot for the *K*_*deg*_, *K*_*up*_, and *K*_*NS*_ parameters in the spleen compartment of the model. The blue line is the model output using optimized parameters obtained via the sensitivity analysis. Other lines show the model output with unoptimized *K*_*deg*_, *K*_*up*_, and *K*_*NS*_ parameters.

Following the sensitivity analysis in model A, model B, and the branching model, values for 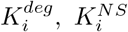, and 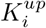 were determined and are available at the SI GitHub link. It is logical to assume that non-specific uptake should be greatest in the liver and gut because they are considered clearance organs. However, the local sensitivity analysis revealed that the highest non-specific uptake rate was in the lungs. This discrepancy is due to the presence of potentially invalid experimental data. It was apparent that no matter how high the liver *K*^*NS*^ value was set during the sensitivity analysis, the biodistribution produced by the model was always significantly smaller than the experimental data set. In fact, forcing a match to the liver data can only be realized at the expense of violating the conservation of mass, which indicates an experimental error in the reporting of the liver data. It is important to note that earlier iterations of the model not described in the paper used other experimental data sets for validation, and the liver biodistribution output matched the experimental data well. So, we conclude that the experimental data reported by [23] for the liver is invalid. As for the gut and other *K*^*NS*^ values, they were entirely arbitrary. Since the experimental data set used for validation did not report NP biodistribution data for gut or other, we could not perform a valid sensitivity analysis on this parameter for these compartments.

### Model Validation

#### Computational Performance Metrics

The unbranched model was solved using MATLAB’s ode15s solver for stiff systems and also using various stiff solvers from Julia’s DifferentialEquations.jl Library. Both A and B models was run from 0 to 10,000 seconds (2.78 hours) to reflect the time scale of a given experimental data set. The branched model was run from 0 to 10,000 seconds on Julia using the same solvers from DifferentialEquations.jl but only from 1 millisecond on MATLAB due to computational constraints. The simulations were performed on a 2020 MacBook Pro with 1.4 GHz Quad-Core Intel core i5 processor and 8 GB 2133 MHz LPDDR3 memory. The timestep of integration was optimized considering stability and the conservation of mass as two metrics. While the stiff solvers in Julia’s DifferentialEquations.jl package do not require a user-defined timestep, for models A and B the timestep of integration, Δ*t*, was 1 millisecond, and was 1 nanosecond for the branched model, in MATLAB. All plots were generated through Julia.

#### Comparison to Experimental Data

To ensure that the model output can be considered accurate, the model was validated using data collected from an experimental study [23]. This study used a variety of sizes of PEGylated gold nanoparticles and sampled the biodistribution at various time points (5, 30, 60, 120 minutes) in five locations in a mouse. Using WebPlotDigitizer (https://apps.automeris.io/wpd/), attenuation values for 100np NP were extracted from Figure 5 (Figure in [23]) and converted to %ID/g values using the relationship presented in Figure 6 (b) of [23]. Then, using given organ weights from [21], %ID/t (percent of injected dose in whole tissue) was calculated. These %ID/t values determined experimentally in [23] were then compared to the model output to ensure the compartmental model is indeed predictive. Single time points for the liver and spleen were reported reliably, so these data sets were used for model validation. While this paper only describes validation with one experimental study, it is important to note that previous iterations of the model used a variety of other experimental studies for validation.

**Fig 5.**
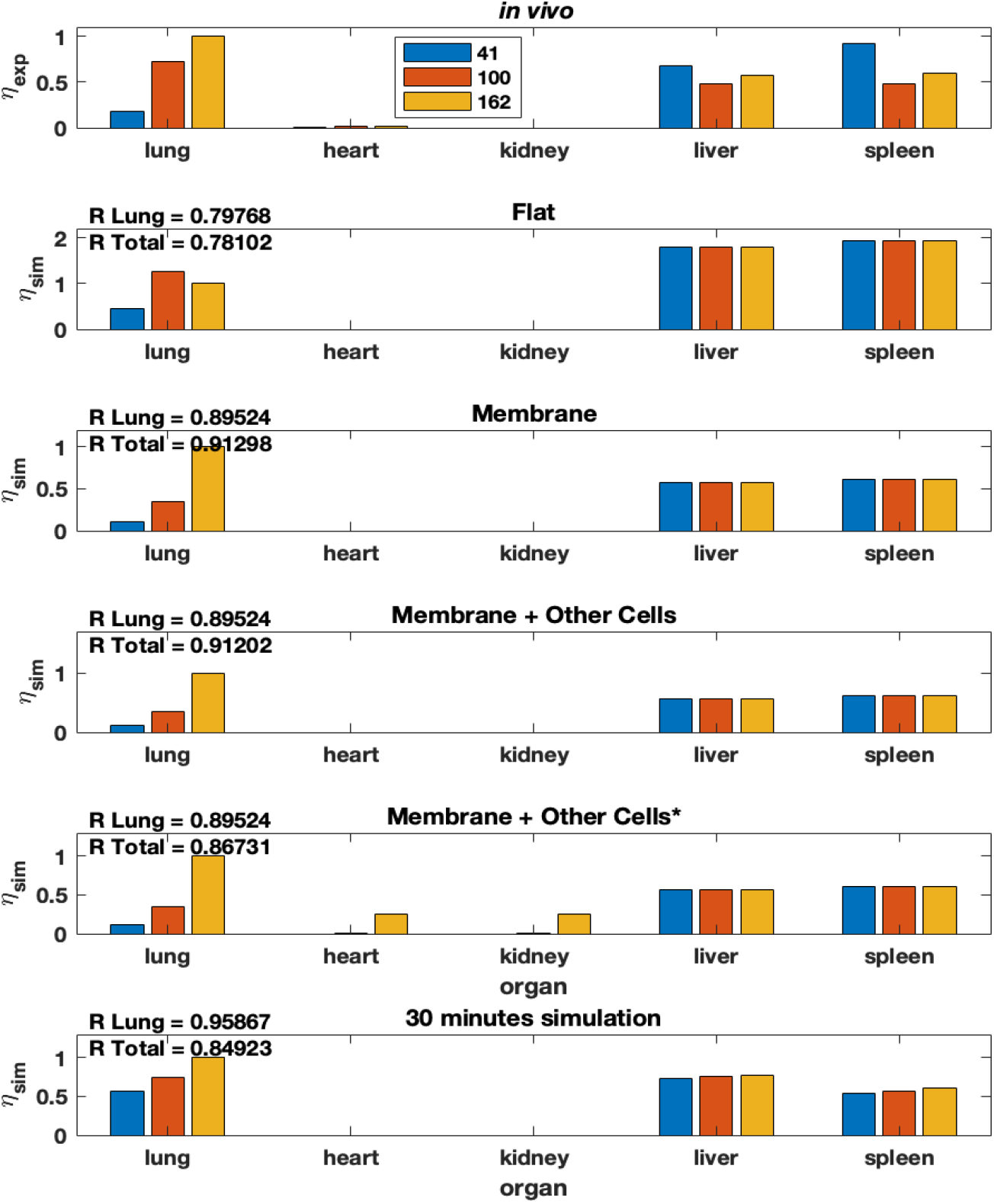
Modified model from Ramakrishnan (2016) [16] that includes non-specific uptake. reported for several membrane conditions, antibody concentrations, and organ combinations.

**Fig 6.**
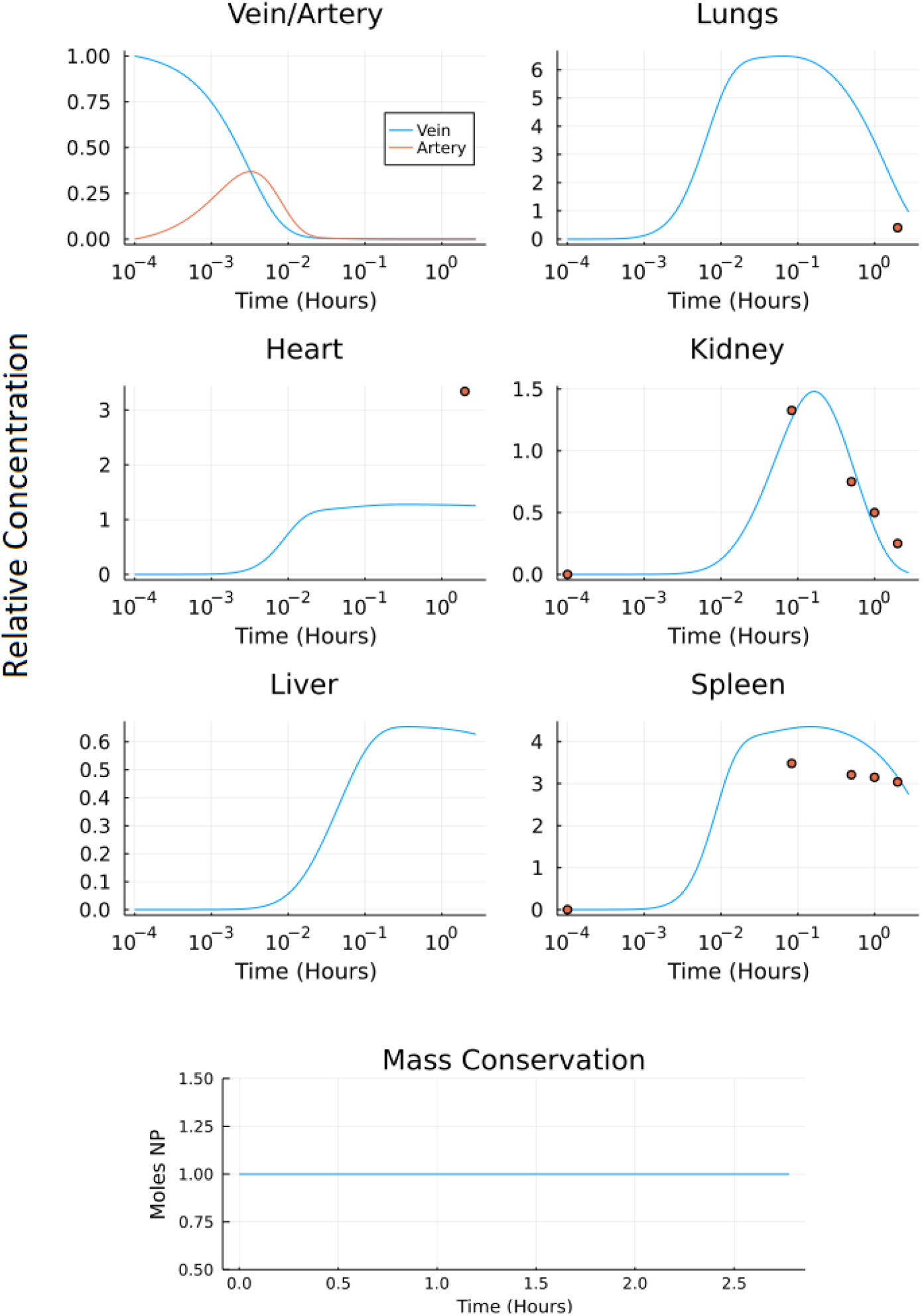
Output of model A: a) the normalized concentration of NP in each of the five organ compartments and the arteries and veins of the original model, and b) the conservation figure.

#### Mass Conservation Analysis

A conservation analysis was employed in every iteration of the model to ensure that each model described can be considered a closed system. The conservation curve of the model is the sum of NP that have been degraded (Equation (14)) as well as those previous time points in every compartment of the model (arterial, venous, organs, vasculature, endothelial cell, and tissue) Equation (15). Mass is conserved in the model when the line formed by all data points from Equation (15) over time *t* has a slope of 0. Equation (16) is used for mass conservation analysis when we setup the system on a molar basis and the mass is conserved where the moles of NPs in all compartments add up to give initial value of NPs in venous compartment.

#### Stiffness of the Unbranched Model

The stiffness ratio characterises the stiffness of the system. For a system of linear ordinary differential equations 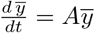, The stiffness ratio is defined as the ratio of the largest and smallest eigenvalue of the matrix A.

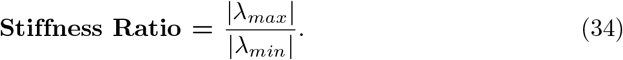

Table 6 in supporting information shows the dependence of stiffness ratio on model parameters, based on the stiffness ratio of our system, we chose ode15s in MATLAB and QNDF, Rodas4, KenCarp4, TRBDF2 and RadauIIA5 from Julia’s DifferentialEquations.jl package to solve our system. All the mentioned solvers are stiff ODE solvers.

### Quasi Steady State Approximation (QSSA)

The system of ODEs described by our PBPK model is a stiff system. The stiffness in the system arises because of the large differences in order of magnitude of model parameters; in our system we have very fast binding and unbinding of NPs to endothelial layer and very slow uptake of NPs into tissue. The model can be made non-stiff by omiting the ODEs containing parameters at both time scales i.e. very fast (*K*_*on*_, *K*_*off*_) and very slow (*K*_*UP*_, *K*_*NS*_). A more qualitative explanation can be given by looking at the results from branched model simulation; it can be seen in the Fig A in supporting information that concentration profiles in the vasculature and endothelial layer quickly drop to zero after the initial spike. Therefore the model can be approximated by omitting the differential equations related to the vasculature and endothelial layer. Consider the two subsystems

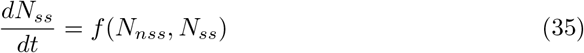

and,

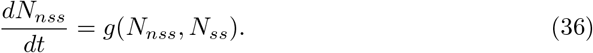

where *N*_*ss*_ denotes the moles of NP particles in vasculature and endothelial in all organs and *N*_*nss*_ denotes the moles of NPs in vein, artery and tissue compartment in all the organs. This reduced system consists of ODEs for concentration profiles in vein, artery and organ tissue (9 in total). The system is much less stiff than the complete systems because *K*_*on*_ and *K*_*off*_ not do occur in the reduced system of ODEs. Functions f and g consists of linear combinations of terms *N*_*ss*_ and *N*_*nss*_ for different organs, because the original system of ODEs is a linear system. Then the QSSA involves setting:

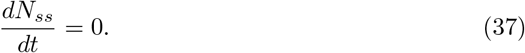

We then use the values of *N*_*ss*_ obtained by solving equation (37) as constants in equation (36), this is no different than solving a system of ODEs in Julia with an extra step of solving a system of linear equations ((37)). Since the reduced system is nonstiff equation (36) can be solved with any nonstiff/explicit ODE solver.

### Neural Networks to Solve ODEs

It has been shown in [25] that Neural Networks can be used to solve partial differential equations; we use the same protocol to solve our system of ODEs using a neural network. Our system being a stiff one makes it ideal for testing the performance of a new method and comparing it against highly optimized ODE solvers. It has been shown [26] that neural networks can be used to solve ODEs when the system of ODEs is nonstiff. We used QSSA to make our PBPK model nonstiff (stiffness ratio ∼10^4^) and trained a neural network to learn the solution to the ODEs.

Consider a system of first-order 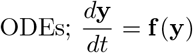, where both **f** and **y** are vectors of same size. Hypothesize that the solution can be approximated using a neural network, with trainable parameters *θ*, i.e.

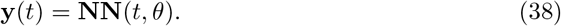

The derivative of the output from the neural network can be approximated by a finite difference scheme or can be obtained using autograd functionality of a neural network. Here we use a first-order finite difference scheme.

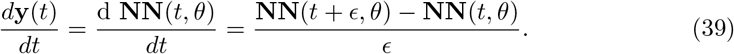

The loss function for this problem is:

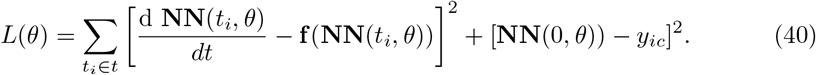

The first term in the loss function is the squared error between LHS and RHS of the ODEs, computed using forward pass through the Neural Networks and the second term is the squared error between true initial condition and predicted initial condition. The solution **NN**(*t, θ*\*) is a unique solution to the system of ODEs, where,

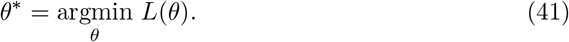

Eq (41) is the optimization problem aiming to minimize the above defined loss in terms of neural network parameters *θ*.

The procedure described in Algorithm 1 aims to solve Eq (35) and (36) iteratively. Notations used in the algorithm mean the same as described in these equations.

## Results

### Biodistribution from Steady State Model Agrees with Murine In Vivo Data

Using the steady-state model equation (Equation (2) in Methods), we plot the %idg values in each of the five organs over three different antibody concentrations on the surface of NP (41, 100, and 162 antibodies coating the surface of the NP), four different theoretical membrane properties (flat, membrane, membrane & present macrophages, membrane & present and active macrophages), and five different organs (lung, heart, kidney, liver, spleen). Additionally, the in-vivo experimental %idg data is plotted, in Figure 5 and R values representing the correlation between the lung %idg values of the given model and experimental lung values, as well as R values that show the correlation between the %idg values of all organs of the model output and the experimental data are given. It is important to note that the total R values across all organs are much higher in the modified model discussed in this paper compared to the original model, comparisons of R values in both models is shown in Table 5 in supporting information. This suggests that the incorporation of non-specific uptake into the model results in a much more physiologically relevant model. We also compare the results obtained from a 30-minute simulation of an unbranched model with experimental data (reported at a 30-minute time-stamp). We observe that by performing local sensitivity analysis for area and volume of the endothelial layer and organ tissue respectively, we obtain a high R for lungs compared to all other models. The excellent agreement of the steady-state data with the 30-minute time data from the transient model suggests that the steady-state model is a good approximation for time scales.

#### Algorithm 1 Physics Informed Neural Network with QSSA

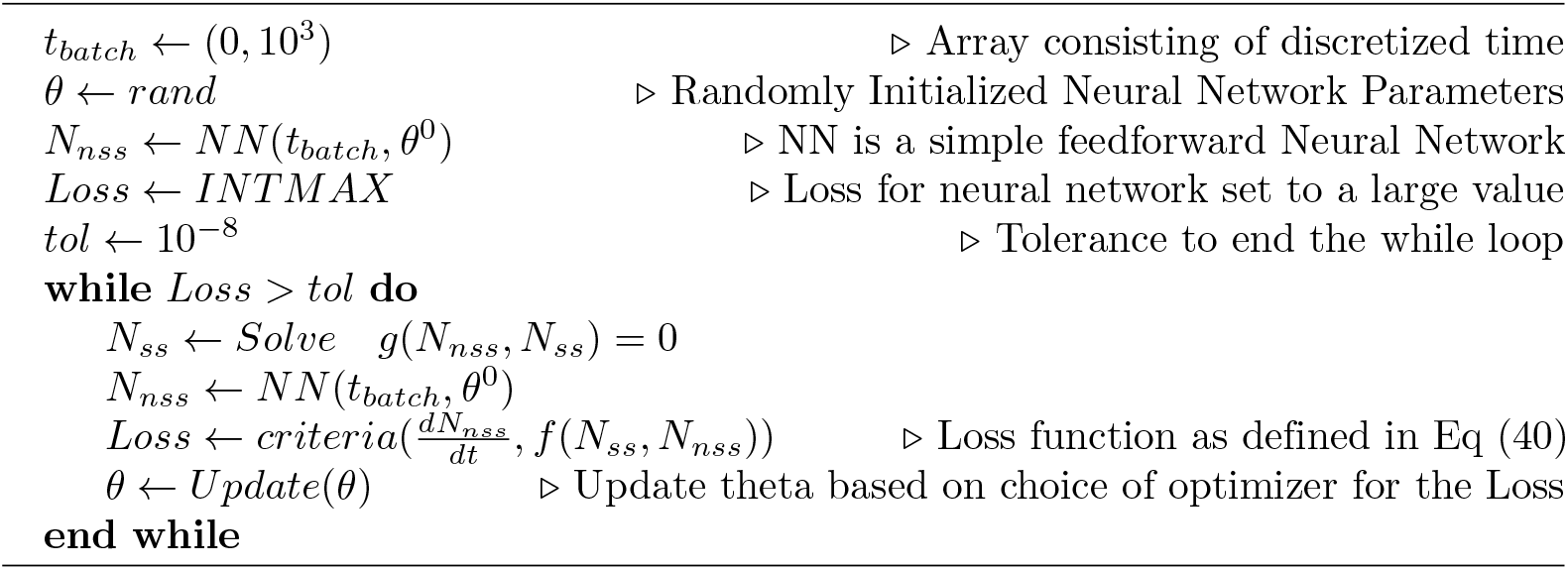

### Compartmental Model B Provides a Better Representation of the Temporal Biodistribution Observed in Experiments

Both compartmental models (A and B) were solved using a variety of stiff solvers available in Julia’s DifferentialEquations.jl package and the MATLAB ode15s solver (for stiff systems) for a time period of 10,000 seconds (2.78 h). The output and corresponding conservation analysis graphs of models A and B are shown in Figures 6 and 7 respectively.

**Fig 7.**
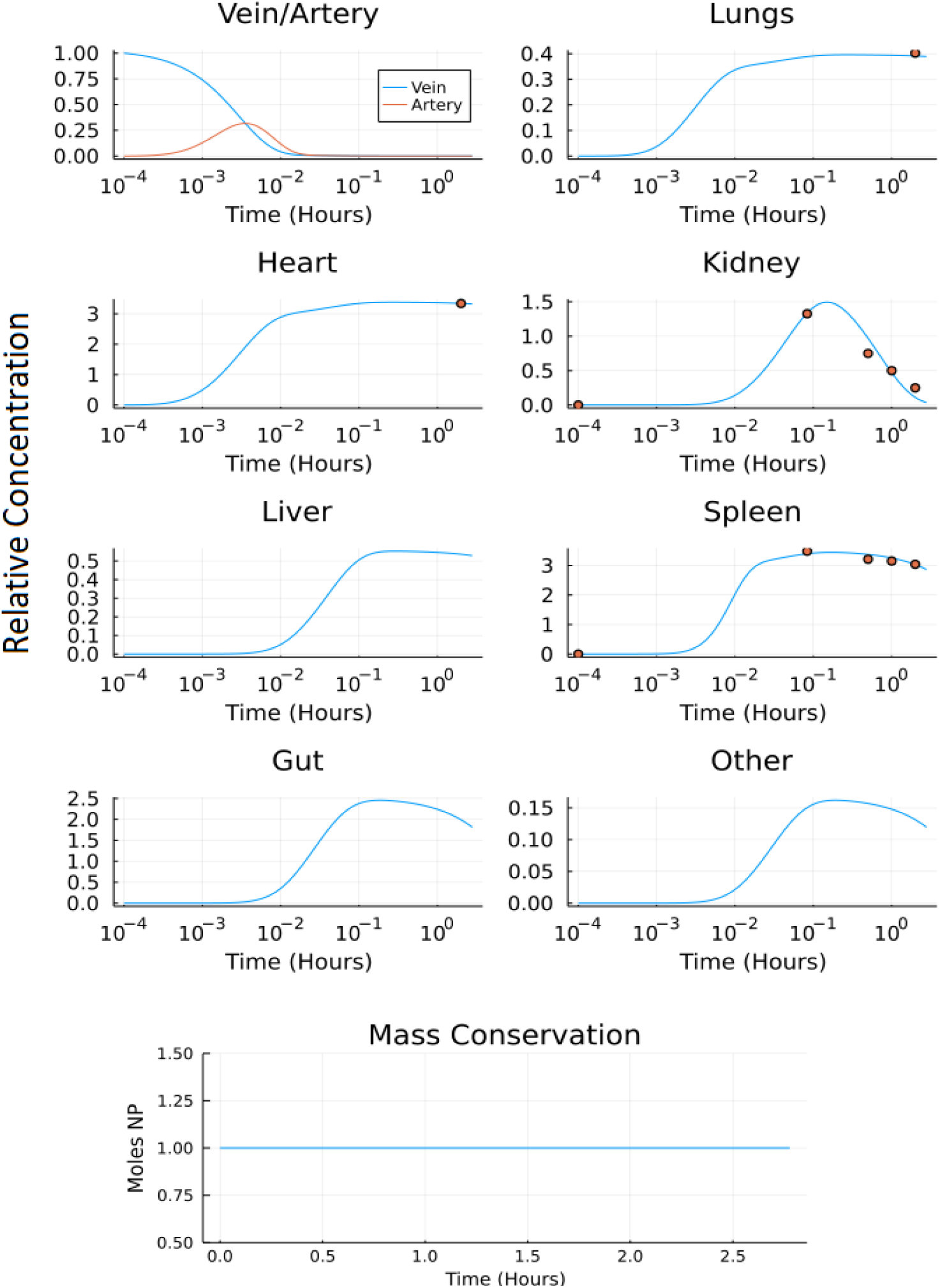
Output of model B: a) the normalized concentration of NP in each of the seven organ compartments and the arteries and veins of the model, and b) the conservation figure.

The normalized root mean squared deviation values (NRMSD) were calculated comparing the model output to the experimental data set where the experimental data set provided time-series data (kidney, liver, spleen). The NRMSD values in model A were 1.0503, 37.4273, and 0.1025 for the kidney, liver, and spleen respectively. The NRMSD values in model B were 0.6735, 20.1909, and 0.0614 for the kidney, liver, and spleen respectively. The NRMSD values were lower in model B than model A suggesting the modified model (model B) more accurately represents the experimental data set [23].

The ODE solvers in Julia’s DifferentialEquations.jl package does not require user-defined timestep and exact mass conservation was obtained by using any of the available stiff solvers. MATLAB’s ode15s solver requires a user-defined timestep (Δ*t*). Timestep was chosen so the system remains stable, and so the system exhibits mass conservation. The Δ*t* of 0.001 seconds was chosen for both compartmental models A and B, for the simulation time of 10,000 seconds. If the Δ*t* was increased beyond 0.001 seconds, the system became unstable and mass was not conserved. On the other hand, if Δ*t* was decreased, the model was unable to complete running due to the computing constraints of MATLAB. It is important to note that mass conservation is only accurate to the order of Δ*t* but integration is valid to a higher order. So, to run the model for a time scale similar to that of experimental studies while maintaining stability and conservation in the system, a Δ*t* of 0.001 for models A and B is necessary.

### Branched Model Predicts a Delayed Temporal Response Compared to the Lumped Compartmental Model

The development of the branched model is incredibly important because it provides a more accurate representation of the circulatory system; specifically, the surface area to volume ratio of the blood vessels in the branched model is more representative of in vivo mouse models. The framework of the branched model allows for the incorporation of more specific hydrodynamic interactions and margination allowing for a physics-based prediction of NP biodistribution, enabling us to account for characteristics such as NP size, shape, and surface chemistry in the model, following the theory in [27]. Additionally, the construction of the branched model will allow for *K*_*on*_ to change depending on vessel diameter and blood flow rate. This cannot be done in models A and B without empirical measurements.

The branched model was solved using Julia’s DifferentialEquation.jl [28] package. Namely, the stiff solvers, QNDF, Rodas4, KenCarp4, TRBDF2 and RadauIIA5 were able to solve for *t* = [0, 10^4^]*s*. Whereas the most efficient stiff solver in MATLAB (ode15s) was unable to solve the system of ODEs. Figure 8 shows the comparison between the molar profiles for the branched and unbranched models.

**Fig 8.**
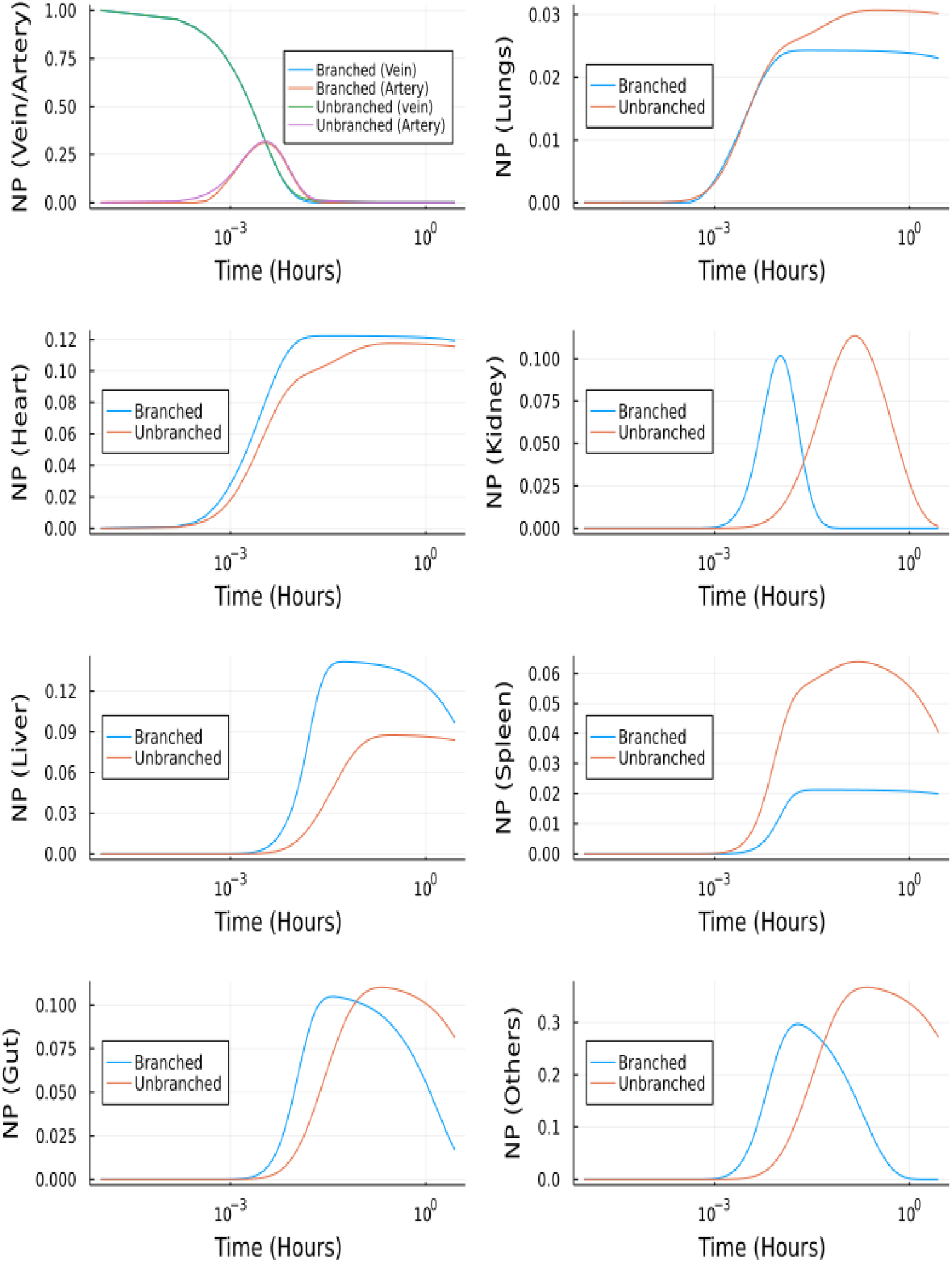
NP concentration vs Time for the branched and unbranched models:. For the organs which have *φ >* 1 the onset is quicker. For lungs *φ* ∼ 1 and for spleen *φ <* 1.

### Effect of Nanoparticle Size on Biodistribution

The branching model construction allows for exploration of the effect of differing NP sizes on biodistribution. The effect of nanoparticle size on biodistribution was also explored using the branched model and nanoparticle diameters of 4nm, 15nm, 50nm, 79nm, and 100nm. In Fig 9, it can be seen that the 5-minute simulation result from our branched model is in good agreement for 5-minute experimental data from [23] for the kidney. Even though we were not able to get an exact match for Liver and Spleen, we observed similar trends between experimental data and simulation data.

**Fig 9.**
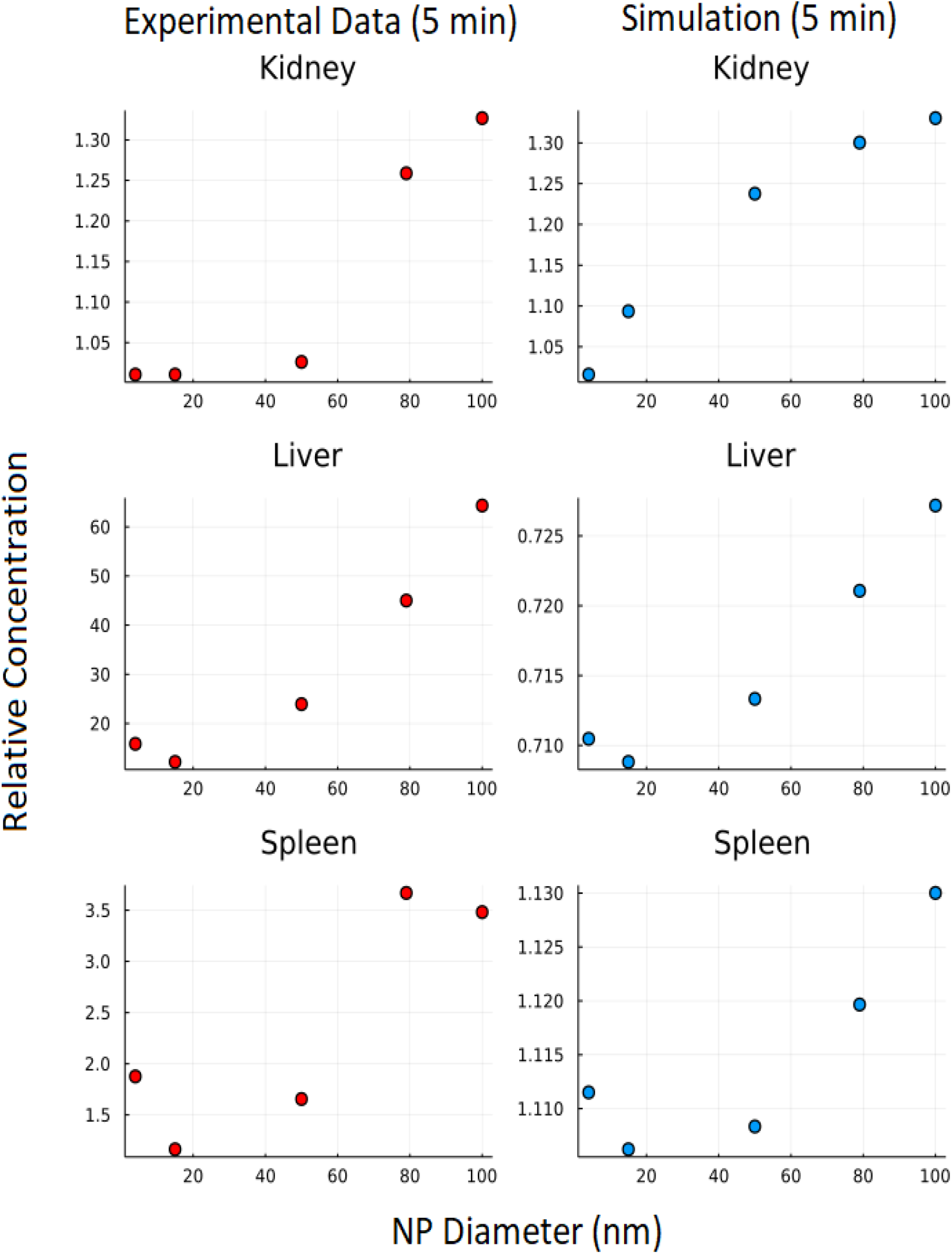
Effect of NP size: Y axis represents relative concentration of nanoparticles at 5 minutes w.r.t initial concentration.

### Model Reduction and Performance of Newer Solvers

#### Quasi Steady State Approximation (QSSA) for the unbranched model

The unbranched model consists of 23 ordinary differential equations, i.e. two equations for vein/artery, seven equations for branched vasculature, seven equations for branched endothelial layer, and seven equations for organ tissues. The reduced system after using QSSA includes only nine equations. The system of linear equations for vasculature and endothelial layer (14 equations) is solved using the similar approach.

Figure 10 depicts the comparison between the unbranched model’s QSSA solution and the unbranched model’s complete solution. The difference in the solutions can be explained based on the vasculature and endothelial profiles for the unbranched model. The difference in profiles for some organs (Kidney, Liver, Gut, Others) is due to the non-zero gradient in profiles of NP bound to endothelial and vasculature for large times, as shown in Fig B in supporting information.

**Fig 10.**
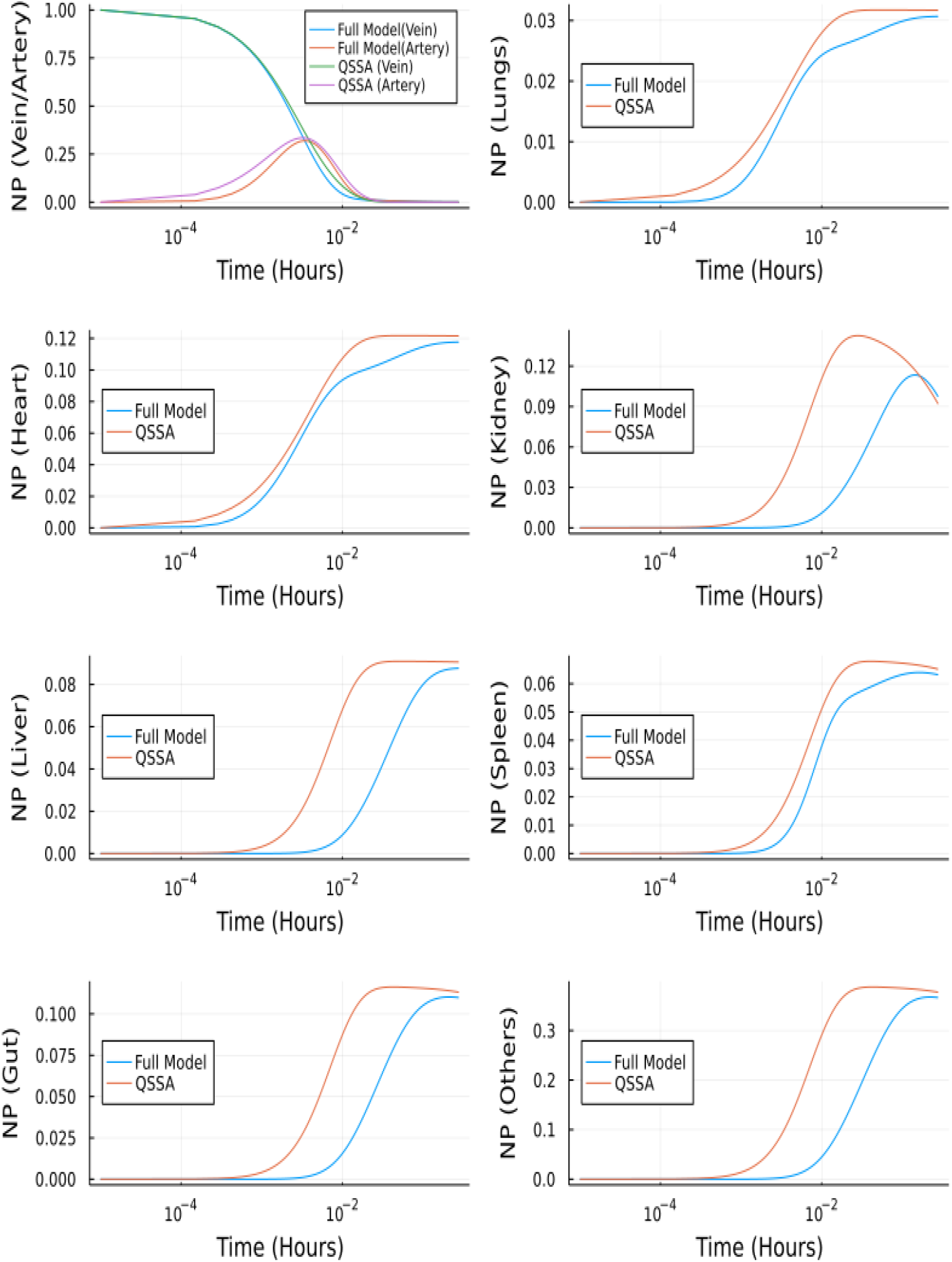
Comparison between complete model and QSSA for unbranched model:

#### QSSA for the branched model

The full branched model consists of 457 ordinary differential equations, i.e. two equations for vein and artery, 32×7 equations for branched vasculature, 32×7 equations for branched endothelial layer, and seven equations for organ tissues. The reduced system after using QSSA has nine equations. The system of linear equations for vasculature and endothelial layer (64*7 equations) is solved using the backslash (*A\b*) operator in Julia, and the values of y are updated in this way at every iteration. However, this makes the task computationally more intensive and takes longer than solving for the complete model, but the objective is to reduce the system’s stiffness. After using QSSA the system of ODE was solved using Tsit5 solver in Julia, which is a nonstiff solver. Figure 11 shows the comparison between QSSA and the complete solution of the branched model. The difference in initial onset can be explained from Figure S1 in supporting information, where we can see the gradient is zero for most of the time but there is a spike initially. QSSA for the branched model is a better approximation compared to QSSA in the unbranched model because of the more rapidly vanishing gradients in the former’s case.

**Fig 11.**
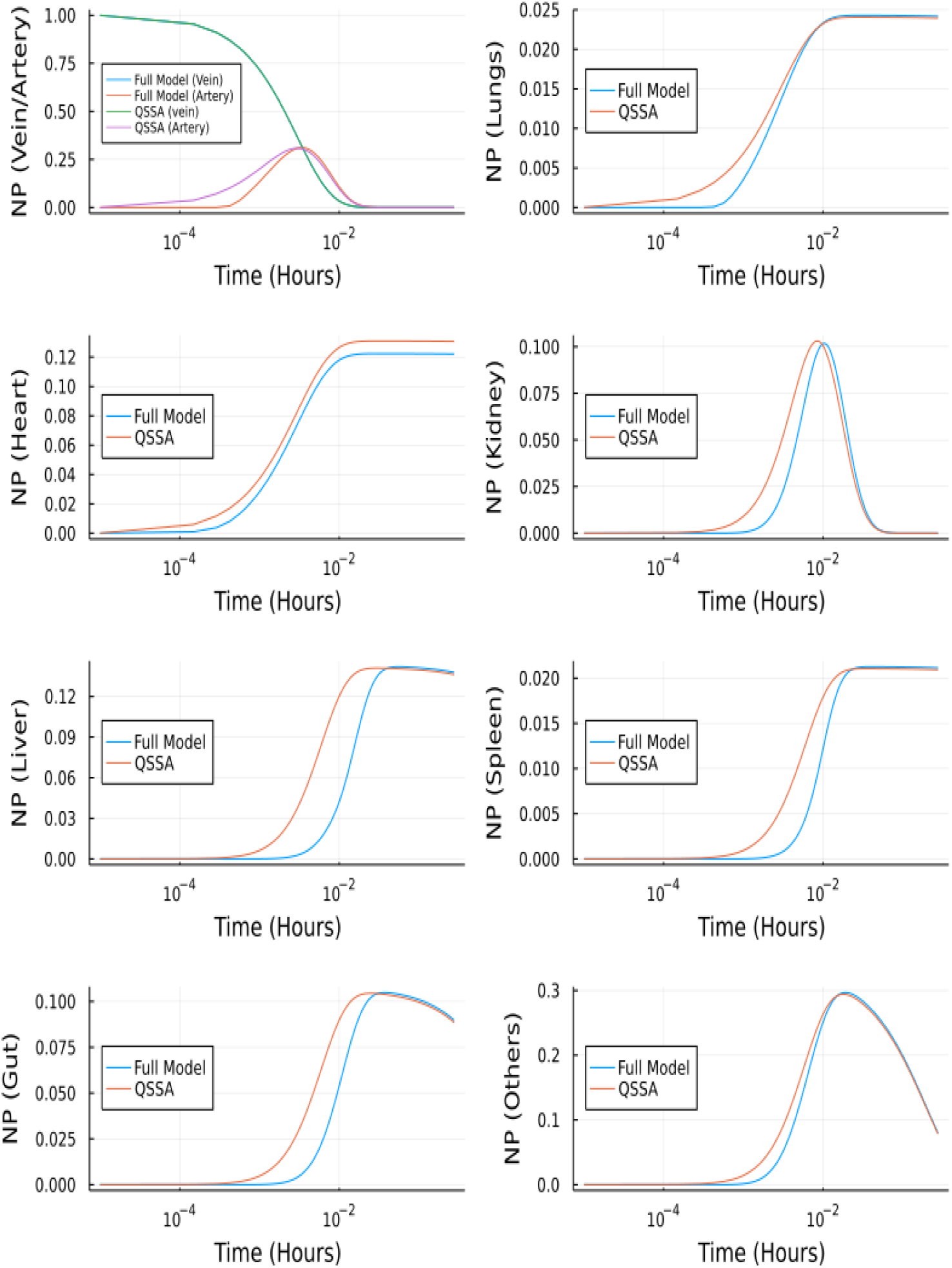
Comparison between complete model and QSSA for branched model.

#### Coupling QSSA and Neural Networks

Using the methods described in the above sections, we solved the concentration profile in the tissue, vein and artery compartments. The neural network predicts these nine outputs, which are used to update the steady-state solution of the remaining 14 equations iteratively.

The neural network was trained for 30000 epochs on a CPU. The neural network architecture consists of an input layer to which discretized time is given as input, two hidden layers each followed by a hyperbolic tangent activation and an output layer consisting of 9 outputs followed by Sigmoid activation. The input to the neural network is a batch of equally spaced numbers between [0, 1], and the first-order derivative of the neural network is scaled with maximum time (*t*_*max*_ = 10^3^) before computing the loss function.

Fig 12 depicts the comparison between the Neural Network solution and the solution obtained using a nonstiff ODE solver (Tsit5 in Julia). This method demonstrates the ability of Neural Networks to solve ODEs. However, we tested this implementation of a simple feedforward neural network to solve the system of ODEs using QSSA. For the full model (without QSSA), we found that the ODE solvers from Julia’s DifferentialEquations.jl library are more adept and a neural network fails to solve the full system.

**Fig 12.**
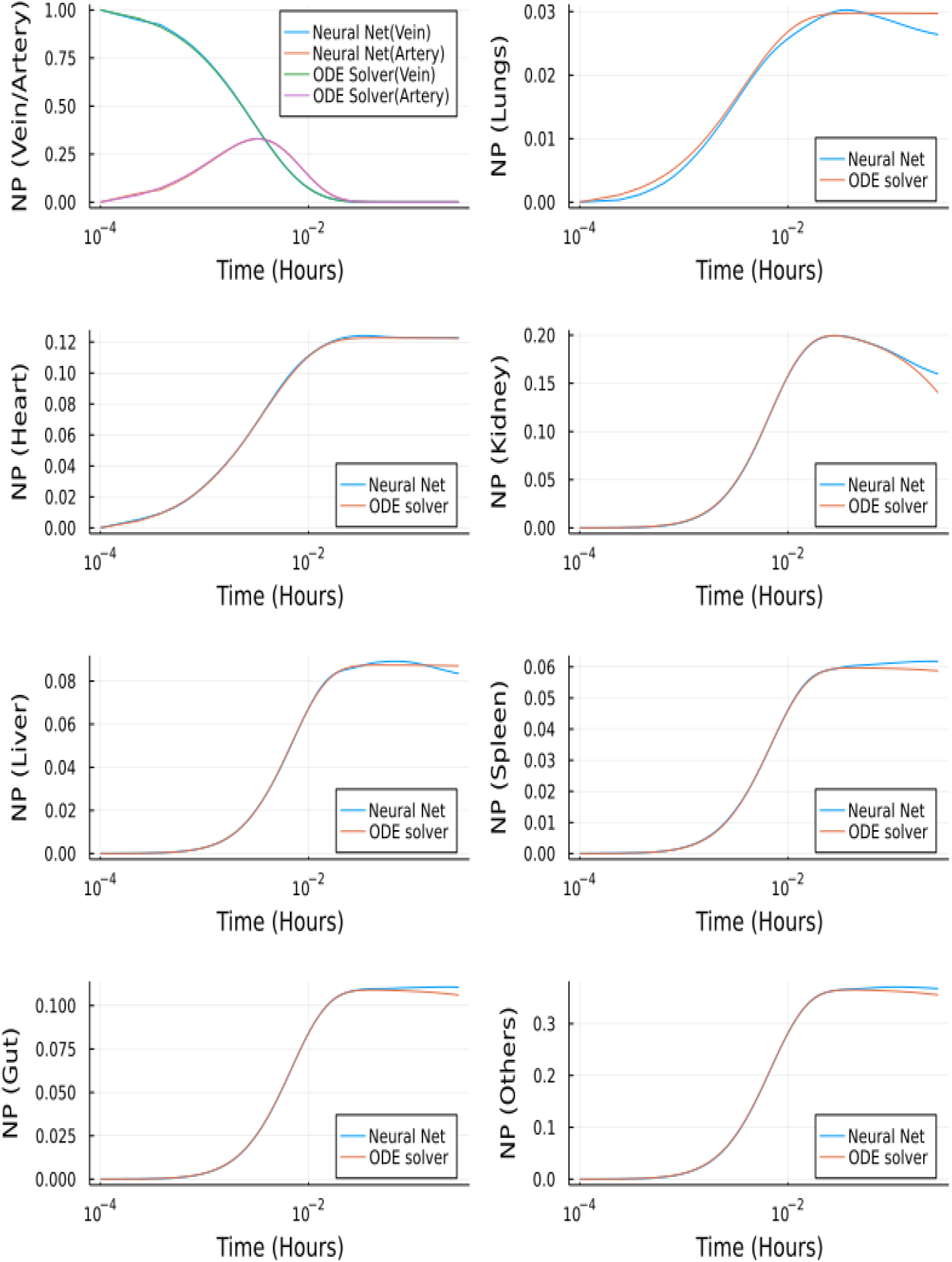
**Comparison** between Neural Networks and Tsit5 with QSSA for unbranched model.

## Discussion

In clinical settings, the use of nanotechnology, including drug-carrying nanoparticles (NP), has increased in recent years. However, the range of NP applications, target, and physical characteristics significantly impede the ability of NP to be researched effectively as bench-to-bedside therapeutics. To this end, researchers have begun to turn toward physiologically based pharmacokinetic (PBPK) models to guide experimentation and better understand the targeting behaviour of various nanoparticle compositions in the human body. A multiscale computationally driven model with physiologically relevant inputs can be utilized to determine organ-specific biodistribution since the physiological and hydrodynamic factors governing NP biodistribution and tissue targeting involve mechanisms that operate at different timescales. NP behaviour must be understood at every level to create a comprehensive multiscale model. This includes the binding landscape of a NP in the presence of an endothelial cell layer. A previous multiscale PBPK model has determined binding constants of intracellular adhesion molecule 1 (ICAM1) coated NPs to endothelial cell surface receptors in mice and humans by utilizing the biophysical properties of the antibody to receptor interactions, and the cell surface [16]. However, the nonspecific uptake, or uptake via passive diffusion in the intercellular cleft, is not accounted for in that model. The purpose of this current study was to 1) modify an existing steady-state PBPK model [16] to incorporate NP uptake via nonspecific transport, 2) develop a novel multiscale PBPK compartmental model to predict temporal effects, and 3) introduce a compartmental branched vascular model that can predict the effect of NP size, 4) perform validation with experimental murine biodistribution data.

The original steady-state model from [16] was modified by adding a nonspecific uptake term that represents NP uptake via passive diffusion through the intracellular cleft. The addition of nonspecific uptake increased the predictive ability of the model, as evidenced by higher R values when compared to an experimental data set than the original model. This suggests that incorporating a nonspecific uptake term into future models is necessary to increase physiological relevance and predictive ability. Next, a novel multiscale PBPK compartmental model was created to predict the continuous temporal biodistribution of NP in five to seven organs. Two versions of this model were created, model A and model B, which differ based on the number of organ compartments represented (model A: 5, model B: 7), and the way flow is routed through the model (Model B is more physiologically relevant). Models A and B were represented with a system of ODEs (Model A: 17 ODEs, Model B: 23 ODEs) solved using stiff solvers from Julia’s DifferentialEquations.jl package and MATLAB’s ode15s solver, then validated with experimental data. The predictive ability of Model B was greater than Model A as evidenced by the normalized root mean squared deviation (NRMSD) analysis. Finally, a branched model was developed to create a more detailed and physiologically relevant version of the basic compartmental model while still maintaining the simplistic compartmental foundation. The branched network consists of a branched vascular tree that begins at the main arteries and veins and bifurcates into the capillary beds, connecting the arterial and venous branching networks. This branching model was represented with 457 ODEs which were solved using Julia. The branching framework allowed for customized output based on NP size. *K*_*on*_ and *K*_*off*_ values can be calculated and are dependent on NP size. Biodistribution for NP size of 4nm, 15nm, 50nm, 79nm, and 100nm was plotted.

The compartmental models’ A and B differed slightly in their composition, with model B more physiologically relevant. Model A consists of five organ compartments (lungs, heart, kidneys, liver, spleen), while model B includes the addition of two additional compartments (gut, other). Additionally, the lung circulation has been separated out to keep track of oxygenation and oxygen distribution, and gut and spleen compartments are coupled to the liver compartment in model B. A local sensitivity (4) analysis was performed on several parameters: *K*_*deg*_, *K*_*up*_, and *K*_*NS*_ to determine the values of these parameters that would best fit the experimental data set available. *K*_*up*_ and *K*_*NS*_ had a similar effect on the model output across organ compartments. When increased, the slope of the biodistribution curve over time would increase, with a higher plateau concentration. When *K*_*deg*_ is increased, the slope of the biodistribution curve over time would decrease, typically decreasing the plateau concentration and also shortening the length of the plateau, leading to a quicker decrease in NP concentration in a given compartment.

The multiscale PBPK model framework presented in this paper presents a significant advance because the predictive ability is purely mechanism-based. Multiscale physics-based modeling allows for the system’s behaviours and interactions to be completely described mathematically, rather than relying on empirical observations and data to make predictions. Typical PBPK models ([8], [10]) are generally empirically based and do not describe the entire behaviour of the system. Creating a purely physics-based PBPK model allows for more customization. For example, in this case, it allows NP composition to be varied by changing certain model parameters to reflect differing NP surface chemistry or size. This is advantageous since NP exist in many forms with various surface chemistries, compositions, and sizes and allows for further model customization in the future.

To continue to evolve this model to allow for customization beyond NP size, additional modules can be added in the future. Incorporating a module that defines internalization rates of varying NP surface chemistries is vital to extending this model to other NP besides ICAM coated. Additionally, hydrodynamic interactions in the bloodstream, immune system effects, and separating various types of NP degradation can be added to the model to increase the physiological relevance and translational potential.

The branching model results in an incredibly large and stiff system of ODEs. To attempt to combat these issues, the stiff MATLAB ode solver, ode15s, was used. MATLAB produced biodistribution graphs, but only from 0-1 milliseconds. This is because a small time-step needed to be used to ensure stability within the model. If the time-step was increased beyond one nanosecond, the system became unstable, and if the run time was increased beyond one millisecond with the one nanosecond time-step, MATLAB became unresponsive. So, it is clear that there are computing power limitations in MATLAB solving large stiff ODE systems. We used stiff equation solvers from DifferentialEquations.jl package in Julia to address this issue. All the stiff solvers from the package successfully solved the system large times.

The physiologically relevant PBPK model can produce a correct output describing the biodistribution of NP. While the neural net ODE solver was successfully demonstrated here, the full power of neural networks can be realized by embedding the multiphysics in the neural networks. Eventually, this computationally driven PBPK model could be used for developing a Neural Network to create a reduced but accurate model for determining NP biodistribution, with more efficiency than the PBPK compartment model. An input to the neural network could be characteristics of the NP such as NP size, vesicle cargo, and concentration of surface proteins. The discrete NP concentrations in each organ as determined by the neural network could be trained against the continuous PBPK compartmental model output (described in this study) at a given time point. Utilizing ML techniques in this model will allow for a much more efficient and automated predictive model for determining NP biodistribution. However, it is necessary to construct an informative PBPK compartmental model to ultimately use in the training process of this Neural Network [25], [26].

## Conclusion

In this study, we successfully advanced existing steady-state multiscale PBPK models [16] to incorporate NP uptake via nonspecific transport, as evidenced by an ability to better predict in vivo biodistribution data (determined by R). Specifically, the total R values across all organs are much higher in the modified model presented in this paper than the original steady-state model.

Secondly, we developed novel multiscale PBPK compartmental models to predict temporal effects of NP distribution. We developed Model A (5 organ compartments, simple configuration) and Model B (7 organ compartments, more physiologically relevant configuration) and solved with both MATLAB ode15s solver and a variety of stiff solvers available in Julia’s DifferentialEquations.jl package, for 10,000 seconds (2.78 h). to determine NP biodistribution temporally, and model outputs were compared to in vivo experimental data. According to NRMSD, Model B more accurately represents the experimental data set used for comparison [23].

Thirdly, we introduced a compartmental branched vascular model that can predict the effect of hydrodynamic interactions that depend on NP size, flow, and vasculature network properties. This branched vascular model is the most physiologically relevant of the models described, as it provides the most accurate representation of a circulatory system. The branched model framework allows for incorporating specific hydrodynamic interactions and marination, which allows a more physics-based prediction of biodistribution, which allows us to consider more specific NP characteristics, like NP size. We explored the effect of NP size (4nm, 15nm, 50nm, 79nm, 100nm) on biodistribution and determined that a five-minute model simulation agrees well with five-minute experimental data.

Finally, we proposed efficient solvers for the stiff systems that make up the models described above and make the solvers compatible with machine learning modules like neural networks that can incorporate multiphysics models into PBPK workflow. The unbranched model can be reduced from 23 equations to 9 equations using QSSA, and the outputs of the reduced model and full model are similar, with differences that can be explained by the non-zero gradient in profiles of NP bound to endothelial and vasculature at large times. The branched model was also reduced from 457 equations to 9 equations using QSSA. Again, the outputs of the reduced and full branched models are similar, and differences in initial onset can be explained by the gradient being zero for most of the time except for an initial spike in the vasculature and endothelial. We concluded that QSSA for the branched model is better than QSSA in the unbranched model due to more rapidly vanishing gradients in the former. Finally, QSSA was coupled to a neural network and determined that the output of the neural network was the same as that of the ODE solver, demonstrating the ability of a neural network to solve ODEs.

This model can continue to evolve to allow for customization beyond NP size by adding modules in future model iterations. For example, incorporating a module that describes internalization rates of varying NP surface chemistries to describe biodistribution of other NP beyond those that are ICAM coated. Additionally, incorporating immune system effects and more specific hydrodynamic interactions into the model would increase NP’s physiological relevance and translational potential to be utilized more frequently in the clinic for targeted therapies.

## Supporting Information

**Supporting Information Figures and Tables** Fig A: Molar profile of NP bound to endothelial layer for branched model. Fig B: Molar profile of NP bound to endothelial layer for unbranched model. Table 1: *K*_*on*_ values for different organs as a function of NP diameter. Table 2: *K*_*off*_ values for different organs as a function of NP diameter. Table 3: log(*K*_*EC*_) values for different organs as a function of NP diameter. Table 4: *K*_*deg*_, *K*_*UP*_, *K*_*NS*_, rates in each organ compartment. Table 5: **R**^2^ value comparison between original Ramakrishnan 2016 model [16] and the modified model. Table 6: Dependence of stiffness ratio on *K*_*on*_

The MATLAB and Julia code (which includes data used to run the models) for this study can be found in the Multiscale PBPK Nanoparticle Biodistrubtion GitHub repository (https://github.com/emmamglass/Multiscale-PBPK-Nanoparticle-Biodistribution-Model)

## Acknowledgments

We thank David Cormode for providing us with experimental validation data, and Greg Conradi Smith for aiding in the continuation of this work at The College of William and Mary.

## Conflict of Interest Statement

The authors declare that the research was conducted in the absence of any commercial or financial relationships that could be construed as a potential conflict of interest.

## Funding

This work was supported by the NSF-funded Science and Technology Center, Center for Engineering Mechanobiology (CEMB), award number CMMI-1548571, by the Penn Engineering Blair Scholar Fellowship Fund, the NIH Physical Sciences in Oncology/ Cancer Systems Biology Consortium Summer Internship, and the William and Mary Charles Center Honors Research Fellow Program. This study has received funding from the National Institutes of Health under U01CA250044.

## Notes

### Competing Interest Statement

The authors have declared no competing interest.

## References

1. Blanco E, Shen H, Ferrari M. Principles of nanoparticle design for overcoming biological barriers to drug delivery. Nature Biotechnology. 2015;33(9):941–951. doi:10.1038/nbt.3330.

2. He H, David A, Chertok B, Cole A, Lee K, Zhang J, et al. Magnetic Nanoparticles for Tumor Imaging and Therapy: A So-Called Theranostic System. Pharmaceutical Research. 2013;30(10):2445–2458. doi:10.1007/s11095-013-0982-y.

3. Anselmo A, Mitragotri S. Nanoparticles in the Clinic. Bioengineering & Translational Medicine. 2016;1. doi:10.1002/btm2.10003.

4. Mitragotri S, Lammers T, Bae YH, Schwendeman S, De Smedt S, Leroux JC, et al. Drug Delivery Research for the Future: Expanding the Nano Horizons and Beyond. Journal of Controlled Release. 2017;246:183–184. doi:https://doi.org/10.1016/j.jconrel.2017.01.011.

5. Hua S, de Matos MBC, Metselaar JM, Storm G. Current Trends and Challenges in the Clinical Translation of Nanoparticulate Nanomedicines: Pathways for Translational Development and Commercialization. Frontiers in Pharmacology. 2018;9:790. doi:10.3389/fphar.2018.00790.

6. Eckmann DM, Bradley RP, Kandy SK, Patil K, Janmey PA, Radhakrishnan R. Multiscale modeling of protein membrane interactions for nanoparticle targeting in drug delivery. Current Opinion in Structural Biology. 2020;64:104–110. doi:https://doi.org/10.1016/j.sbi.2020.06.023.

7. Radhakrishnan, Ravi and Yu, Hsiu-Yu and Eckmann, David M. and Ayyaswamy, Portonovo S. Computational Models for Nanoscale Fluid Dynamics and Transport Inspired by Nonequilibrium Thermodynamics. Journal of Heat Transfer. 2016;139:033001. doi:https://doi.org/10.1115/1.4035006.

8. Li M, Al-Jamal KT, Kostarelos K, Reineke J. Physiologically Based Pharmacokinetic Modeling of Nanoparticles. ACS Nano. 2010;4(11):6303–6317. doi:10.1021/nn1018818.

9. Lin Z, Monteiro-Riviere NA, Riviere JE. A physiologically based pharmacokinetic model for polyethylene glycol-coated gold nanoparticles of different sizes in adult mice. Nanotoxicology. 2016;10(2):162–172. doi:10.3109/17435390.2015.1027314.

10. Yuan D, He H, Wu Y, Fan J, Cao Y. Physiologically Based Pharmacokinetic Modeling of Nanoparticles. Journal of Pharmaceutical Sciences. 2019;108(1):58–72. doi:https://doi.org/10.1016/j.xphs.2018.10.037.

11. Rao GG, Landersdorfer CB. Antibiotic pharmacokinetic/pharmacodynamic modelling: MIC, pharmacodynamic indices and beyond. International Journal of Antimicrobial Agents. 2021;58.

12. Snoek E, Jacqmin P, Sargentini-Maier ML, Stockis A. Modeling and simulation of intravenous levetiracetam pharmacokinetic profiles in children to evaluate dose adaptation rules. Epilepsy Research. 2007;76.

13. Sharma RP, Schuhmacher M, Kumar V. The development of a pregnancy PBPK Model for Bisphenol A and its evaluation with the available biomonitoring data. Science of the total environment. 2018;624.

14. Husoy T, Martinez MA, Sharma RP, Kumar V, Andreassen M, Sakhi AK, et al. Comparison of aggregated exposure to di(2-ethylhexyl) phthalate from diet and personal care products with urinary concentrations of metabolites using a PBPK model – Results from the Norwegian biomonitoring study in EuroMix. Food and Chemical Toxicology. 2020;143.

15. Li M, Nguyen L, Subramaniyan B, Bio M, Peer CJ, Kindrick J, et al. PBPK modeling-based optimization of site-specific chemo-photodynamic therapy with far-red light-activatable paclitaxel prodrug. Journal of Controlled Release. 2019;308.

16. Ramakrishnan N, Tourdot R, Eckmann D, Ayyaswamy P, Muzykantov V, Radhakrishnan R. Biophysically inspired model for functionalized nanocarrier adhesion to cell surface: roles of protein expression and mechanical factors. Royal Society Open Science. 2016;3.

17. Ayyaswamy PS, Muzykantov V, Eckmann DM, Radhakrishnan R. Nanocarrier Hydrodynamics and Binding in Targeted Drug Delivery: Challenges in Numerical Modeling and Experimental Validation. Journal of Nanotechnology in Engineering and Medicine. 2013;4(1). doi:10.1115/1.4024004.

18. Radhakrishnan R. A survey of multiscale modeling: Foundations, historical milestones, current status, and future prospects. AIChE Journal. 2020;67. doi:10.1002/aic.17026.

19. Agrawal N, Radhakrishnan R. The Role of Glycocalyx in Nanocarrier-Cell Adhesion Investigated Using a Thermodynamic Model and Monte Carlo Simulations. The journal of physical chemistry C, Nanomaterials and interfaces. 2007;111:15848–15856. doi:10.1021/jp074514x.

20. Yang J, Wang Y. Design of vascular networks: A mathematical model approach. International journal for numerical methods in biomedical engineering. 2013;29. doi:10.1002/cnm.2534.

21. Diehl L, Morse M. A Comparison of Selected Organ Weights and Clinical Pathology Parameters in Male and Female CD-1 and CByB6F1 Hybrid Mice 12-14 Weeks in Age. Charles River, Spencerville. 2015;.

22. Boswell CA, Mundo EE, Ulufatu S, Bumbaca D, Cahaya HS, Majidy N, et al. Comparative Physiology of Mice and Rats: Radiometric Measurement of Vascular Parameters in Rodent Tissues. Molecular Pharmaceutics. 2014;11(5):1591–1598. doi:10.1021/mp400748t.

23. Dong Y, Hajfathalian M, Maidment P, Hsu J, Naha P, Si-Mohamed S, et al. Effect of Gold Nanoparticle Size on Their Properties as Contrast Agents for Computed Tomography. Scientific Reports. 2019;9. doi:10.1038/s41598-019-50332-8.

24. McKenzie M, Ha S, Rammohan A, Radhakrishnan R, Natesan R. Multivalent Binding of a Ligand-Coated Particle: Role of Shape, Size, and Ligand Heterogeneity. Biophysical Journal. 2018;114:1830–1846. doi:10.1016/j.bpj.2018.03.007.

25. Raissi M, Perdikaris P, Karniadakis GE. Physics-informed neural networks: A deep learning framework for solving forward and inverse problems involving nonlinear partial differential equations. Journal of Computational Physics. 2019;378:686–707. doi:https://doi.org/10.1016/j.jcp.2018.10.045.

26. Ji W, Qiu W, Shi Z, Pan S, Deng S. Stiff-PINN: Physics-Informed Neural Network for Stiff Chemical Kinetics. The Journal of Physical Chemistry A. 2021;125(36):8098–8106. doi:10.1021/acs.jpca.1c05102.

27. Jabeen Z, Yu HY, Eckmann DM, Ayyaswamy PS, Radhakrishnan R. Rheology of colloidal suspensions in confined flow: Treatment of hydrodynamic interactions in particle-based simulations inspired by dynamical density functional theory. Phys Rev E. 2018;98:042602. doi:10.1103/PhysRevE.98.042602.

28. Rackauckas C, Nie Q. Differentialequations.jl–a performant and feature-rich ecosystem for solving differential equations in julia. Journal of Open Research Software. 2017;5(1).

29. Innes M, Saba E, Fischer K, Gandhi D, Rudilosso MC, Joy NM, et al. Fashionable Modelling with Flux. CoRR. 2018;abs/1811.01457.

